# Principles and functional consequences of plasmid chromatinization in mammalian cells

**DOI:** 10.1101/2025.05.27.656122

**Authors:** Benjamin J. Mallory, Thomas W. Tullius, Carina G. Biar, Jonas A. Gustafson, Stephanie C. Bohaczuk, Danilo Dubocanin, Lea M. Starita, Andrew B. Stergachis

**Affiliations:** Department of Genome Sciences, University of Washington School of Medicine, Seattle, WA, USA; Brotman Baty Institute for Precision Medicine, Seattle, WA, USA; Lewis-Sigler Institute for Integrative Genomics, Princeton University, Princeton, NJ, 08544; Division of Genetic Medicine, Department of Pediatrics, University of Washington, Seattle, Washington 98195, USA; Molecular and Cellular Biology Program, University of Washington, Seattle, Washington 98195, USA; Division of Medical Genetics, University of Washington School of Medicine, Seattle, WA, USA; Department of Genetics, School of Medicine, Stanford University, Stanford, CA, 94304

## Abstract

Plasmids have fundamentally transformed how we resolve regulatory grammar across the tree of life. However, although chromatin plays an integral role in regulating the function of regulatory elements along the nuclear genome, our understanding of how, or whether, similar chromatin structures form on plasmids transfected into mammalian cells remains limited. We demonstrate that plasmid single-molecule chromatin fiber sequencing (plasmid Fiber-seq) can accurately map chromatin architectures along individual, full-length transfected plasmid molecules at near single-nucleotide resolution. Application of plasmid Fiber-seq to diverse plasmids and cell lines demonstrates that plasmids are chromatinized in a sequence-dependent organized manner and adopt a heterogeneous and incomplete chromatin architecture relative to nuclear-encoded chromatin fibers. We show that the focal occupancy of nucleosomes and transcription factors along transfected plasmids is central to their transcriptional activity within mammalian cells, and demonstrate that plasmids indeed are capable of recapitulating nuclear genome-encoded chromatin architectures, although not always. Finally, we demonstrate that combining plasmid Fiber-seq with high-throughput reporter assays can establish the molecular mechanisms underlying pathogenic non-coding variants, including disentangling the effects of transcriptional activators and repressors with near-single-nucleotide resolution. Overall, our findings reveal the principles by which plasmid-based assays can be used for accurate fine-scale mapping of chromatin-dependent regulatory grammar.

## Main Text

Plasmids, first characterized as any extrachromosomal genetic element in the early 1950’s^1^, have fundamentally transformed molecular biology and served as the foundation for core technologies in biology ranging from recombinant DNA^2^ to CRISPR genome editing^3^ and therapeutic gene delivery systems^4^. Furthermore, plasmids have proven to be a powerful tool for systematically identifying regulatory elements and quantifying how genetic variants modulate their function through luciferase assays^5^, Massively Parallel Reporter Assays (MPRA)^6^ and Self-Transcribing Active Regulatory Region Sequencing (STARR-seq) assays^7^, among others. Central to these research and clinical applications of plasmids is that when plasmids are transfected into mammalian cells, their encoded regulatory elements are capable of engaging with the mammalian transcriptional machinery. Methods such as luciferase assays, MPRAs and STARR-seq have taken advantage of this feature to functionally characterize millions of regulatory sequences across diverse cell types, deepening our understanding of regulatory grammar. However, these methods rely on the assumption that regulatory elements function similarly when present within the plasmid or endogenous nuclear genome context, an assumption that has previously been questioned^8^.

In the endogenous nuclear genome context, chromatin plays an integral role in regulating the function of regulatory elements, yet our understanding of how, or whether, similar chromatin structures form on plasmids transfected into mammalian cells remains limited. In general, it is well established that plasmids can be occupied by histones when transfected into mammalian cells, as first demonstrated in the 1970s with the viral SV40 plasmid genome^1^ and later shown using non-viral plasmids^9^. Subsequent work using micrococcal nuclease (MNase) digestion and Southern hybridization demonstrated that some plasmids adopt a chromatin pattern characterized by greater spacing between individual nucleosomes compared to the nuclear genome^10^. Modern next-generation sequencing-based chromatin mapping technologies reliant on short-read sequencing^11,12^ are poorly suited for studying how individual plasmids are chromatinized. Specifically, individual cells often contain 100s of plasmids that may be adopting a distinct chromatin architecture, thereby limiting even the ability of single-cell based approaches for uniquely mapping the architecture of a plasmid^13^. Furthermore, existing short-read based methods cannot reliably distinguish whether a read is arising from the plasmid or host genome when the plasmid contains endogenously-encoded regulatory sequences due to mapping ambiguities, thereby preventing the precise delineation of how regulatory elements are chromatinized when present within the plasmid or endogenous nuclear genome.

To address this challenge we developed a plasmid single-molecule chromatin fiber sequencing method (plasmid Fiber-seq) that leverages recent advances in single-molecule chromatin fiber sequencing^14^ to accurately map chromatin architectures along individual, full-length transfected plasmid molecules at near single-nucleotide resolution. Using plasmid Fiber-seq, we dissect the principles underlying plasmid chromatinization, and demonstrate how plasmid chromatinization can affect the outcome of reporter assays and be used to mechanistically resolve the functional impact of pathogenic variants.

## Single-molecule chromatin architectures of transfected plasmids

To study plasmid chromatinization within mammalian cells, we developed plasmid Fiber-seq, which leverages single-molecule long-read chromatin fiber sequencing (Fiber-seq)^14^ to stencil the chromatin architecture along individual transfected plasmids via a nonspecific DNA N^6^-adenine methyltransferase (m6A-MTase). Importantly, as DNA-m6A is effectively not endogenous in multicellular organisms^15,16^, m6A-marked stencils can accurately resolve single-molecule chromatin architectures along individual plasmids with near nucleotide resolution. However, as bacteria often endogenously express m6A-MTases, plasmid Fiber-seq leverages an E. coli strain (ER2796) defective in all major DNA m6A-MTases^17^ as the primary plasmid source prior to mammalian cell transfection^18^ (**Extended Data** Fig. 1). After transfection, isolated nuclei are treated with the non-specific m6A-MTase Hia5^19^, and plasmid DNA is then enriched from total nuclear DNA, linearized using a m6A-insensitive restriction enzyme, and sequenced using highly accurate single-molecule long-read sequencing (**Fig. 1a**).

**Figure 1.**
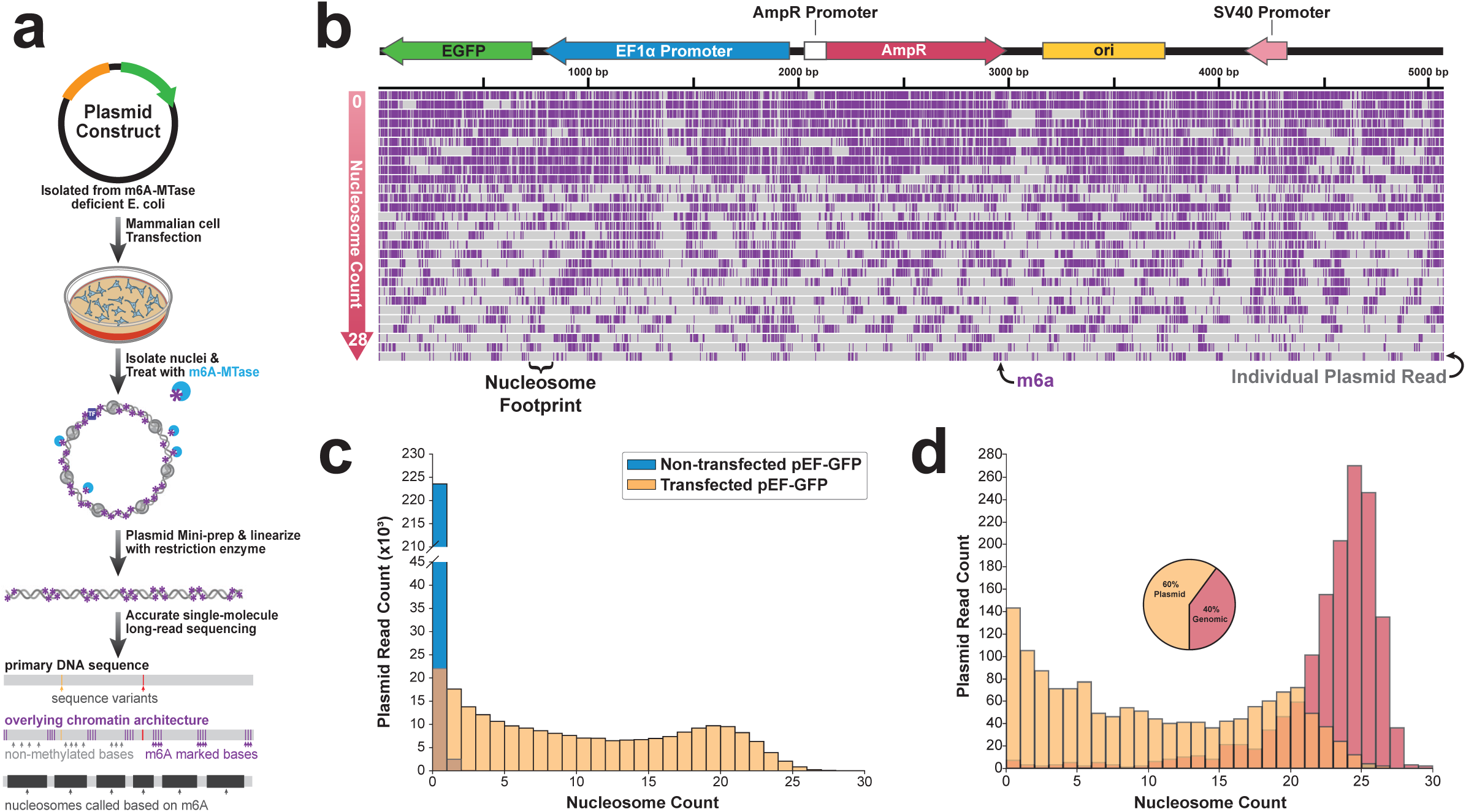
| Plasmid single-molecule chromatin fiber sequencing (plasmid Fiber-seq). **(a)** Schematic for plasmid Fiber-seq using the non-specific adenine methyltransferase Hia5 to stencil single-molecule protein occupancy using m6A. **(b)** Single-molecule full length plasmid Fiber-seq reads of the linearized pEF-GFP construct, demonstrating the heterogeneity of plasmid methylation patterns. Grey bars represent individual full-length plasmid reads and purple dashes denote m6a, marking accessibility. **(c)** Histogram quantifying the number of reads existing in each chromatin state for both non-transfected pEF-GFP and pEF-GFP transfected into HEK293 cells. **(d)** Histogram comparing the quantification of plasmid and genomic reads existing in each chromatin state. Plasmid and genomic reads contain similar length (+/-50 bp) and regulatory sequence context (Methods). Pie chart displays the proportion of total reads from a single plasmid Fiber-seq reaction mapping to either unique plasmid constructs or the hg38 genome.

To test this approach, we transfected HEK293 cells with the commonly used plasmid pEF-GFP^20^, which contains *GFP* regulated by the human EF1α promoter, and performed plasmid Fiber-seq 48 hr post-transfection. Plasmid Fiber-seq resulted in ∼60% of all sequencing reads arising from full-length plasmid constructs (426,622 full-length plasmid reads), with the remaining reads largely arising from randomly sheared genomic DNA. We next applied *fibertools*^18^ and *FiberHMM*^21^ to identify single-molecule putative protein occupancy footprints along each read based on the m6A stencil patterns (**Fig. 1b**). This exposed that the predominant footprint size along individual plasmid fibers was ∼130 bp (**Extended Data** Fig. 2), which is comparable with the average size of nucleosome footprints seen along the sheared nuclear genomic fibers sequenced during the same reaction. Importantly, this footprint size was not seen along untransfected plasmids treated *in vitro* with Hia5 (**Fig. 1c**), indicating that plasmid Fiber-seq can accurately resolve nucleosome architectures along individual transfected plasmids.

## Plasmids adopt a heterogeneous and incomplete chromatin architecture

To determine whether nucleosomes occupy plasmids to a similar degree as they do nuclear genome-encoded chromatin fibers, we first identified sequencing reads that arose from the nuclear genome and were of similar size and regulatory element content as the pEF-GFP sequencing reads from the same plasmid Fiber-seq reaction (Methods). This revealed a tight and right-shifted distribution of nuclear-encoded chromatin fibers, which most frequently contained 24 nucleosome-sized footprints per read (NPR) (**Fig. 1d**). In contrast, 38% of reads from the full-length pEF-GFP plasmid showed fewer than 5 NPR (i.e. <20% of the expected density of nuclear-encoded chromatin fibers -‘lowly chromatinized’) while 26% had 15 or more NPR (i.e., >60% of the density of the expected nuclear-encoded chromatin fibers - ‘highly chromatinized’) (Methods). Among highly chromatinized plasmids, the most frequent state contained 19 NPR, indicating that the pEF-GFP plasmid adopts a heterogenous and incomplete chromatin architecture in HEK293 cells. To further explore this finding, we applied plasmid Fiber-seq to HEK293 cells transfected for 48 hours with an equimolar amount of five distinct plasmids that contained a shared prokaryotic backbone and diverse reporter genes and regulatory elements (**Fig. 2a**). This revealed that whereas nuclear genome-encoded chromatin fibers contained a nucleosome-sized footprint every ∼200 bp (**Fig. 2b**), consistent with prior reports^22^, transfected full-length plasmid fibers contained a nucleosome-sized footprint every ∼286 bp (**Fig. 2b**), demonstrating that plasmids consistently adopt an incomplete chromatin architecture relative to nuclear-encoded chromatin fibers. Notably, the five plasmids differed in the degree to which they adopted a heterogeneous chromatin architecture, with the smallest (2,682 bp) and largest (14,689 bp) plasmids harboring markedly fewer lowly chromatinized reads (6.5% and 1.6%, respectively) relative to the three intermediately sized constructs (4,723 bp, 5,058 bp, and 6,466 bp, with 35%, 28.2%, and 22.8%, respectively) (**Fig. 2c**). Together, this demonstrates that plasmids adopt a heterogeneous and incomplete chromatin architecture characterized by less densely organized nucleosomes compared to nuclear-encoded chromatin fibers.

**Figure 2.**
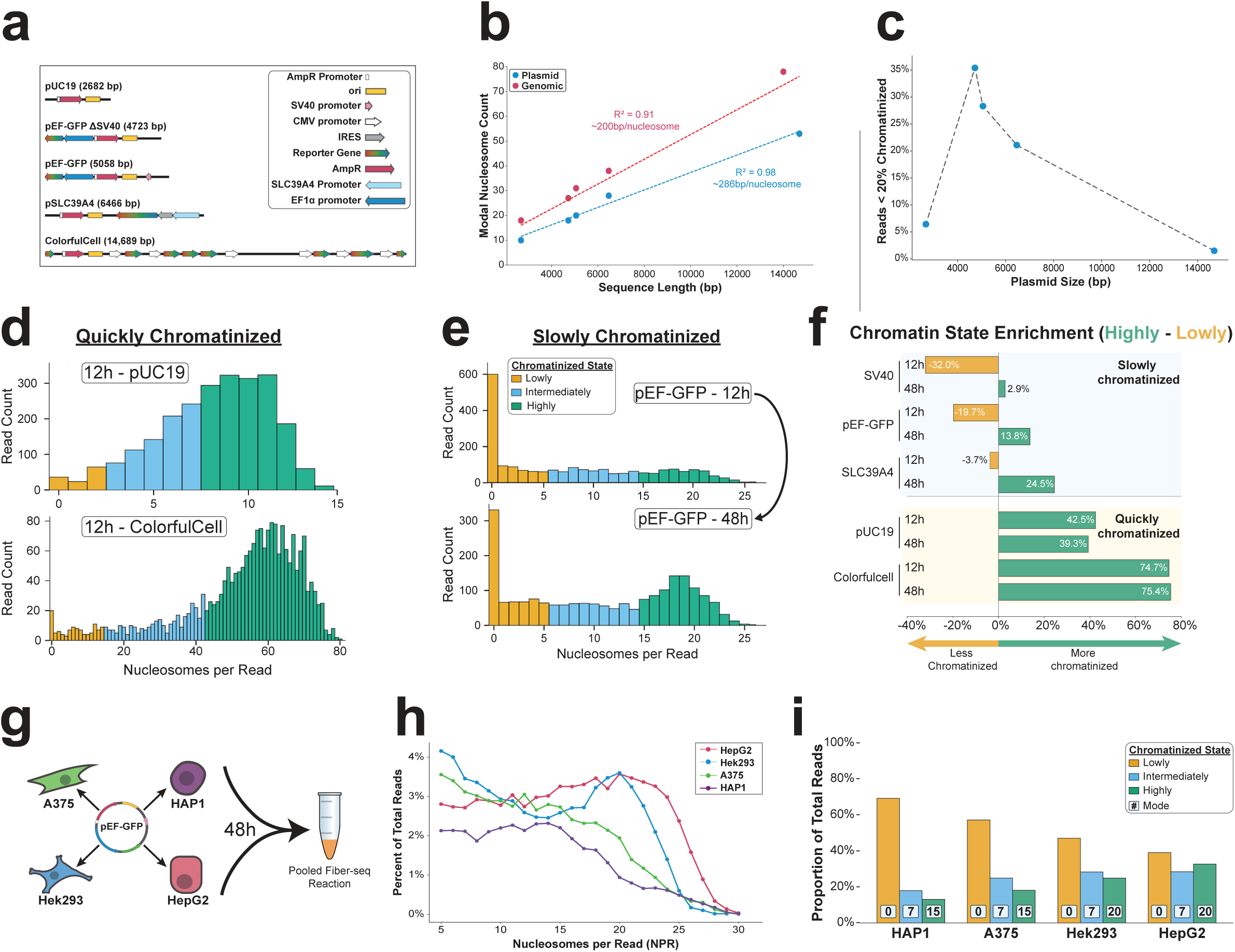
| **Experimental variables alter the chromatin state of transfected plasmids. (a)** Schematic of experimental plasmid constructs and their corresponding sequence elements **(b)** Scatter plot showing the relationship between sequence length and modal nucleosome count for plasmid and genomic reads from the same Fiber-seq reaction, highlighting the linear relationship between read length and nucleosome count, and the decreased density of nucleosomes along plasmid fibers. The modal nucleosome count for plasmid reads is for reads in the highly chromatinized state (i.e. >60% chromatinized (Methods)). **(c)** Scatter plot showing the relationship between sequence length and the proportion of reads existing in the lowly chromatinized state (<20% chromatinized (Methods)). **(d)** Histograms showing the chromatin distribution of plasmid constructs that are quickly chromatinized, reaching a nucleosome-saturated state within 12h following transfection in HEK293 cells. Bin color corresponds the category of chromatin state according to the legend in e. **(e)** Histograms showing the chromatin distribution of plasmid constructs that are slowly chromatinized, with the proportion of reads in the highly chromatinized state gradually increasing from 12 to 48h post-transfection in HEK293 cells. **(f)** Quantification of the difference in the proportion of reads existing in the highly vs lowly chromatinized states at 12h and 48h time points for slowly and quickly chromatinized plasmid constructs transfected in HEK293 cells. **(g)** Diagram demonstrating the experimental approach for assessing the chromatin architecture of pEF-GFP transfected in four different cell types. Nuclei from all cell lines were pooled 48h after transfection and a single plasmid Fiber-seq reaction was performed. **(h)** Line plot showing the chromatin distribution of pEF-GFP in four different cell-types for intermediately and highly chromatinized reads (lowly chromatinzed reads are excluded for visual clarity, but are quantified in i). **(i)** Quantification of the proportion of pEF-GFP reads in each chromatin state across the four different cell types. Bin colors correspond to the chromatin state, with the boxed numbers representing the modal nucleosome count for that state.

To investigate whether the observed heterogeneous and incomplete chromatin patterns vary with time, we transfected the five plasmid constructs into HEK293 cells and performed plasmid Fiber-seq at 12-, 18-, 24-, and 48-hours post-transfection (**Fig. 2d-f**). Notably, we observed that for three of the plasmids, the proportion of plasmids adopting a lowly chromatinized state reduced as a function of incubation time (a decrease from 12h to 48h of 19.5%, 17.3%, and 14.6% for the 4,723 bp, 5,058 bp, and 6,466 bp constructs, respectively) (**Fig. 2e,f**). In contrast, the smallest and largest plasmids, which had the fewest lowly chromatinized reads at 48-hours, showed no appreciable change in the proportion of plasmids adopting a lowly chromatinized state as a function of incubation time (**Fig. 2d,f**). Importantly, all five plasmids lack the ability to replicate within mammalian cells, indicating that plasmid chromatinization can be at saturation in as little as 12 hours, and that the ratio of observed heterogeneous and incomplete plasmid chromatinization patterns progress towards saturation with time.

Transcription-dependent DNA supercoiling is known to disrupt chromatin structure along the nuclear genome^23^, so it is possible that the heterogeneous and incomplete chromatin architectures observed along plasmids could simply reflect the vestigial effects of supercoiled plasmids. To evaluate this, we nicked pEF-GFP and re-ligated it to create a relaxed circular state (**Extended Data** Fig. 3a). We then co-transfected HEK293 cells with an equimolar mix of barcoded relaxed and supercoiled pEF-GFP and performed plasmid Fiber-seq 48 hours post-transfection. We found that supercoiling had only a modest inhibitory effect on plasmid chromatinization, with supercoiled reads showing a ∼7% enrichment of lowly chromatinized reads relative to reads in the relaxed state (**Extended Data** Fig. 3b). However, the relaxed plasmids still adopted a heterogeneous and incomplete chromatin architecture, demonstrating that this is an inherent feature of plasmid chromatinization.

## Determinants of plasmid chromatinization in mammalian cells

Previous studies have shown that the nuclear localization of plasmid DNA typically occurs passively during nuclear envelope breakdown and reformation at telophase^24^. To determine whether this may be driving our observations of heterogeneous and incomplete plasmid chromatinization, we transfected barcoded pEF-GFP plasmids into four mammalian cell lines that markedly differ in their rate of cell division (HAP1 (myelogenous leukemia): doubling time 12-48h, A375 (melanoma): doubling time 6-22h, HEK293 (embryonic kidney): doubling time 18-36h, and HepG2 (liver carcinoma): doubling time 48h) and performed plasmid Fiber-seq on this pooled sample 48h post-transfection (**Fig. 2g**). This revealed that the distribution of chromatinized plasmids markedly diverges between different mammalian cell types in a manner that cannot be fully explained by cell division alone. If cell division were the primary mechanism driving plasmid chromatinization, we would expect cell lines with faster growth rates (HAP1, A375, and HEK293) to show the highest proportion of highly chromatinized plasmids. However, we observed that HepG2, which has the slowest growth rate, actually showed the highest proportion of highly chromatinized plasmids. Further, the NPR mode for the highly chromatinized reads ranged from 20 NPR in HepG2 cells to 15 NPR in HAP1 cells (**Fig. 2h,i**), indicating that plasmids adopt a heterogeneous and incomplete chromatin architecture in a cell-type specific manner that is largely independent of cell division.

In addition, previous studies have shown that specific DNA nuclear targeting sequences can actively drive plasmid nuclear import in non-dividing cells^25–27^, with the SV40 promoter sequence being one of the first described. To assess whether the SV40 promoter influences the rate of plasmid chromatinization, we transfected HEK293 cells with an equimolar mix of pEF-GFP with and without the SV40 promoter sequence and performed plasmid Fiber-seq at 12-, 18-, 24- and 48-hours post-transfection. We observed that the SV40-containing construct consistently had 2-fold higher nuclear read recovery across all time points (**Extended Data** Fig. 4a). Additionally, the SV40-containing construct exhibited a substantially higher proportion of reads in the highly chromatinized state at 12h, and maintained relatively lower proportions of lowly chromatinized reads across all time points (**Extended Data** Fig. 4b). Overall, these results show that the SV40 promoter sequence significantly enhances both nuclear import and chromatinization of plasmid DNA.

## Transfected plasmids are chromatinized in an organized manner

We next evaluated whether chromatin patterns occur in an organized manner along transfected plasmids. Specifically, we used FiberHMM to identify nucleosome-sized and sub-nucleosome-sized footprints along plasmid molecules, and then aggregated these patterns based on each molecule’s overall level of plasmid chromatinization (**Fig. 3a**). We performed this on the pEF-GFP 48-hours post-transfection HEK293 plasmid Fiber-seq sample that was deeply sequenced (3,049,719 reads) to enable evaluation of chromatin features along rare transition states between lowly and highly chromatinized plasmids. Overall, this revealed that the positioning of nucleosome-sized footprints along pEF-GFP is not random, but rather lowly chromatinized reads are preferentially occupied by nucleosome-sized footprints at 6 focal sites along the plasmid, and that nucleosome-sized footprints consistently occupy these 6 sites irrespective of a plasmid’s total chromatinization level (**Extended Data** Fig. 5). Notably, lowly chromatinized reads are also preferentially occupied by sub-nucleosome-sized footprints selectively at these same 6 sites, but the occupancy of these sub-nucleosome-sized footprints dissipates on plasmids that are intermediately or highly chromatinized, opposite to what is seen for nucleosome-sized footprints at these sites.

**Figure 3.**
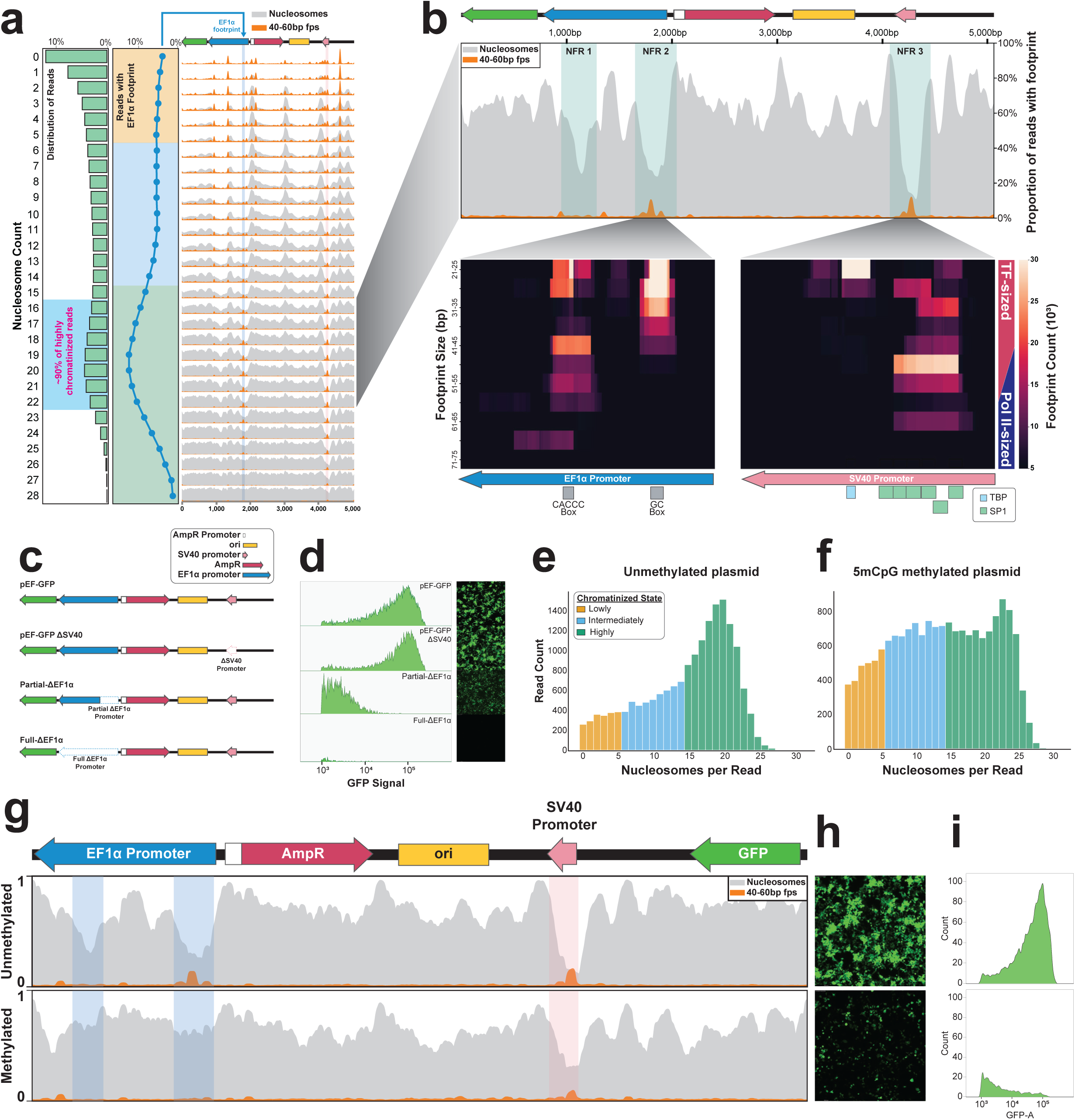
| Transfected plasmids are chromatinized in an organized manner. **(a)** Ridgeplot showing the proportion of reads containing either nucleosome footprints (grey) or 40-60 bp footprints (fps, orange) along each position of the linearized pEF-GFP plasmid. The top schematic outlines the pEF-GFP construct (sequence elements defined in the legend of c). Y-axis for each plot ranges from 0-100% read occupancy, with 40-60 bp footprints scaled-up 4x for visibility. The blue line plot (left) shows the proportion of reads containing a 40-60 bp footprint within the nucleosome free region (NFR 2 in b) of the EF1α promoter across chromatin states, highlighting the relationship of this footprint with plasmid nucleosome occupancy. Colors correspond to lowly (yellow), intermediately (blue) and highly (green) chromatinized states. The grey histogram (left) shows the quantification of reads existing in each chromatin state. **(b)** A smoothed line plot showing the average occupancy of nucleosomes (grey) and 40-60 bp footprints (fps) for pEF-GFP reads containing 16-22 nucleosomes per read. The top schematic represents a map of the linearized pEF-GFP construct (sequence elements defined in the legend of c). Blue boxes highlight the three nucleosome-free regions (NFR) present at these chromatin states. Heatmaps show the diversity of 20-70 bp footprint sizes existing along the sequence corresponding to the location of the 40-60 bp fps peaks present in the EF1α and SV40 promoters, with footprint sizes binned in 5 bp increments. Underlying diagrams highlight the location of the regulatory sequences and corresponding binding motifs. The rectangle (right) highlights the expected size range for transcription factor sized footprints (TF-sized, red) and Pol II sized footprints (Pol II-sized, blue). **(c)** Diagram showing the pEF-GFP deletion constructs generated to validate that NFR associated 40-60 bp footprints correspond to transcriptional activity. **(d)** Quantification of GFP expression as measured by flow cytometry analysis for the corresponding WT or deletion pEF-GFP constructs. **(e)** Histogram showing the chromatin distribution of unmethylated (5mCpG) pEF-GFP 48h post-transfection in HEK293 cells. **(f)** Histogram showing the chromatin distribution of 5mCpG methylated pEF-GFP 48h post-transfection in HEK293 cells. **(g)** Smoothed line plots showing the proportion of reads occupied by nucleosomes (grey) or 40-60 bp footprints (orange) for unmethylated (top) and 5mCpG methylated (bottom) pEF-GFP reads 48h post-transfection in HEK293 cells. Blue boxes highlight the loss of the 40-60 bp footprint and NFRs within the EF1α promoter, and the pink box highlights the decrease in 40-60 bp footprints and NFR within the SV40 promoter. **(h)** Fluorescent image of HEK293 cells transfected with unmethylated or 5mCpG methylated pEF-GFP 48h post-transfection, showing the visual decrease in fluorescence of the 5mCpG methylated plasmid **(i)** Quantification of GFP expression from HEK293 cells transfected with unmethylated or 5mCpG methylated pEF-GFP 48h post-transfection.

To determine whether these 6 sites may be sequence dependent, we performed this analysis on a separate deeply sequenced plasmid Fiber-seq sample (605,714 reads) generated using a 6.4kb pSLC39A4 plasmid that shares an identical 1,937 bp backbone to the pEF-GFP plasmid, but otherwise contains distinct sequence. The pSLC39A4 plasmid similarly demonstrated that lowly chromatinized reads are preferentially occupied by nucleosome-sized footprints at focal sites along the plasmid. Furthermore, the shared backbone present in both pEF-GFP and pSLC39A4 showed a nearly identical pattern of nucleosome- and sub-nucleosome-sized footprints along lowly chromatinized reads (**Extended Data** Fig. 6). Together, this demonstrates that a handful of sites along plasmids are being preferentially occupied by protein complexes that nucleate nucleosome occupancy along transfected plasmids in a largely sequence-specific manner.

## Focal chromatin patterns regulate plasmid function within mammalian cells

We next evaluated whether focal chromatin patterns along highly chromatinized plasmids are related to their transcriptional activity. First, we observed that highly chromatinized pEF-GFP plasmids contained three preferential nucleosome free regions (NFR), one overlapping the SV40 promoter, and two overlapping the EF1α promoter (**Fig. 3b)**. Both the SV40 promoter (NFR3) and upstream EF1α promoter (NFR2) were also preferentially occupied by sub-nucleosome-sized footprints. However, these two sub-nucleosome-sized footprints were retained and even increased along highly chromatinized plasmids, indicating that they have a distinct role in the function of chromatinized plasmids relative to the sub-nucleosome-sized footprints seen at nucleosome nucleation sites.

The single-molecule footprints within the SV40 and upstream EF1α promoter NFRs largely overlapped annotated TF binding elements. Specifically, the SV40 NFR was enriched for 21-30 bp footprints overlapping multiple SP1 and TBP binding elements, as well as a large 46-50 bp footprint that appeared to result from synchronous single-molecule occupancy of adjacent SP1 binding elements. In contrast, the upstream EF1α promoter NFR contained a focal 21-45 bp footprint overlapping a GC box, and a set of discrete footprints overlapping a CACCC box, that ranged from 21-70 bp in length. Notably, the larger 40-70 bp upstream EF1α promoter NFR footprint overlaps the expected size of RNA polymerase II (Pol II)^21^, suggesting that this specific EF1α promoter NFR may be driving GFP expression from the pEF-GFP plasmid.

To test the hypothesis that the upstream EF1α promoter NFR is driving GFP expression, we generated pEF-GFP constructs that contained either a targeted deletion of the upstream NFR (Partial-ΔEF1α), or a complete deletion of the entire promoter (Full-ΔEF1α) (**Fig. 3c**). We observed that the Full-ΔEF1α plasmid resulted in the complete loss of GFP expression, with no new NFRs forming upstream of *GFP* along this construct. In contrast, the Partial-ΔEF1α plasmid resulted in only the loss of the upstream EF1α promoter NFR **(Extended Data** Fig. 7). However, despite the preservation of the downstream EF1α promoter NFR on this plasmid, GFP expression was substantially diminished (**Fig. 3d**), indicating that the upstream EF1α promoter NFR plays a dominant role in driving GFP expression from the pEF-GFP plasmid.

To further disentangle whether these EF1α promoter NFRs are related to pEF-GFP transcription levels, we performed plasmid Fiber-seq on pEF-GFP constructs with or without *in vitro* treatment with the CpG Methyltransferase M.SssI (**Extended Data** Fig. 8), as the CpG methylated plasmid is associated with significantly reduced GFP expression compared to the non-methylated control (**Fig. 3h,i**). Analysis of the corresponding plasmid Fiber-seq data showed that CpG methylation results in an increased rate of lowly chromatinized reads (**Fig. 3e,f**). Furthermore, CpG methylation results in the disappearance of both EF1α promoter NFRs, as well as the Pol II-sized footprints within the upstream EF1α promoter NFR (**Fig. 3g)**. In contrast, the SV40 promoter NFR and sub-nucleosome-sized footprints were still retained, albeit to a lesser degree, along the CpG methylated plasmid. Together, these findings demonstrate that focal nucleosome and TF occupancy patterns along plasmids regulate their transcriptional activity within mammalian cells.

## Plasmids can accurately recapitulate nuclear genome-encoded chromatin architectures

Plasmid-based reporter assays have been foundational to our understanding of gene regulation since they were first introduced in the early 1980’s^28^. However, it remains largely unclear how faithfully nuclear chromatin architectures are recapitulated along plasmid constructs. To resolve this, we used plasmid Fiber-seq, which is grounded in long-read single-molecule sequencing, thereby permitting us to uniquely map chromatin patterns along regulatory elements that are present within both the plasmid and nuclear genome context in the same cell. Specifically, we separately cloned nine extended promoter regions from the HEK293 genome (size range 3,676 to 4,572 bp) into a common plasmid backbone. Three of these promoter regions showed inaccessible chromatin in HEK293 cells, whereas the other six contained between 1 and 5 DNaseI hypersensitive sites within HEK293 cells (**Fig. 4, Extended Data** Fig. 9,10). Plasmid Fiber-seq was performed on HEK293 cells co-transfected with all 9 plasmids, and in parallel we performed Fiber-seq on HEK293 cells, enabling us to directly compare chromatin patterns along the same sequence within a plasmid and endogenous genomic context.

**Figure 4.**
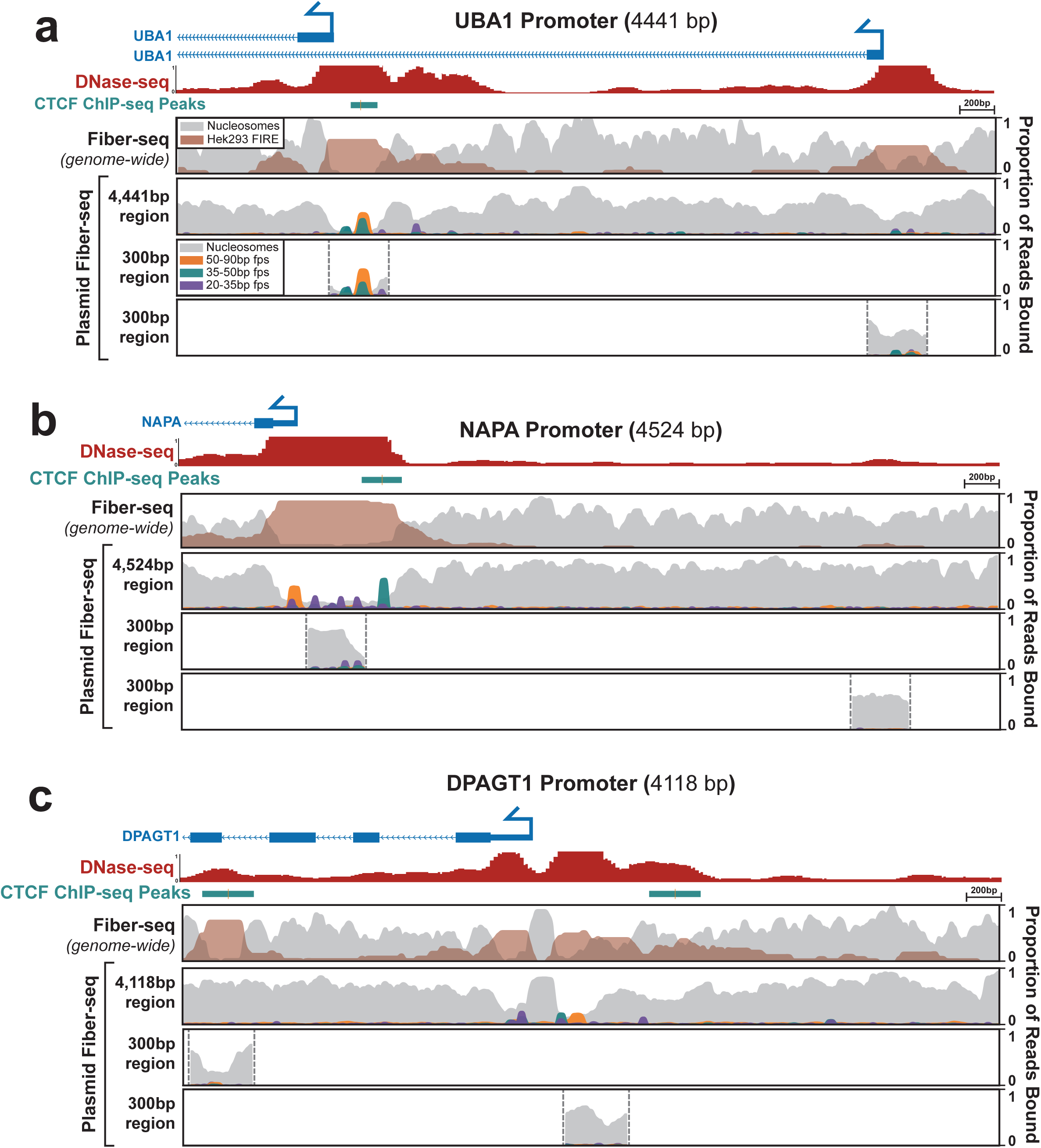
| Comparison of regulatory element chromatin architectures. Comparison of chromatin architectures of the *UBA1* **(a)**, *NAPA* **(b)**, and *DPAGT1* **(c)** promoters. The top panel of each promoter shows the endogenous genomic chromatin architecture for a 4-4.5kb region of promoter as measured by Fiber-seq, with the second panel showing the chromatin architecture of the same sequence within a plasmid context, measured by plasmid Fiber-seq. The bottom two panels show the chromatin architecture of plasmid constructs containing 300 bp promoter fragments centered over the Fiber-seq Inferred Regulatory Element (FIRE) peaks derived from the genomic Fiber-seq data. Top tracks show CTCF ChIP-seq and DNase-seq peaks from Hek293 cells, along with a map of the gene bodies.

Overall, we found that the endogenous chromatin architectures were faithfully replicated on 7 of the nine extended promoter regions. For example, the *MYH7*, *MYBPC3*, and *HBB* promoters are all endogenously inaccessible in HEK293 cells, and we similarly observed no NFRs within these extended promoter regions in the plasmid contexts (**Extended Data** Fig. 9). Similarly, the endogenous *FLCN*, *NAPA*, *UBA1,* and *WASF1* promoters all contain at least one accessible element in HEK293 cells, and we observed that sequence corresponding to these accessible elements preferentially harbored NFRs when present within the plasmid context (**Fig. 4a,b,c, Extended Data** Fig. 10). Furthermore, these NFRs harbored small protein footprints that precisely correspond to well defined TF binding elements, such as the CTCF binding elements in the *NAPA* and *UBA1* promoters. In contrast, the extended *MYC* and *DPAGT1* promoter regions showed clear focal differences in their chromatinization when present within the nuclear or plasmid context. Specifically, although the *DPAGT1* plasmid recapitulated the chromatin architecture of the *DPAGT1* promoter itself, an element in the *DPAGT1* gene body that is highly accessible in the genomic locus was completely inaccessible in the plasmid context. Similarly, although the *MYC* plasmid recapitulated the chromatin architecture of the *MYC* promoter and an upstream CTCF binding element, an intervening element that is highly accessible in the genomic locus was completely inaccessible in the plasmid context.

The above findings demonstrate that plasmids are capable of accurately recapitulating nuclear genome-encoded chromatin architectures. However, these all used 3-5 kb sequences, whereas most plasmids-based reporter assays typically use only ∼300 bp sequences due to the size limitations of array synthesized DNA fragments. To explore how using a smaller sequence context would impact the chromatinization of regulatory DNA within a plasmid construct, we performed plasmid Fiber-seq on 19 constructs that each contained only a 300 bp fragment of the larger 3-5 kb endogenous promoter regions, centered on FIRE peak called from whole-genome Fiber-seq of HEK293 cells. Overall, we found that 14 of the 19 300 bp fragments largely recapitulated the chromatin patterns along the endogenous locus (**Fig.4, extended Data Fig. 9,10**). Specifically, the fragments from the endogenously inaccessible *MYH7*, *MYBPC3*, and *HBB* promoters similarly lacked NFRs when present in the plasmid contexts. Similarly, the *UBA1* and *WASF1* promoters and the *MYC* promoter upstream CTCF binding element showed NFRs and small footprint occupancy patterns along the 300 bp fragments that mirrored those seen on the extended promoter plasmids and that of the endogenous locus.

In contrast, although the extended *FLCN* promoter plasmid construct showed two clear NFRs that mirrored accessible elements present along the endogenous promoter, when these two sites were present as only 300 bp fragments neither one appeared to harbor an NFR. Similarly, although the extended *NAPA* promoter plasmid construct showed a clear NFR that mirrored the accessible promoter element along the endogenous promoter, when only the central 300 bp of this promoter was present within the plasmid context this NFR largely disappeared. Further, whereas the element in the *DPAGT1* gene body was completely inaccessible in the extended plasmid context, the plasmid containing a 300 bp fragment of this element did harbor an NFR that recapitulated endogenous chromatin accessibility. Overall, these findings demonstrate that plasmids are indeed capable of accurately recapitulating nuclear genome-encoded chromatin architectures. However, the surrounding sequence context of the regulatory element plays an integral role in modulating this process which can differ between shorter or longer sequence contexts. Importantly, plasmid Fiber-seq can readily detect differences in chromatin architecture between shorter and longer sequence contexts, thereby informing the experimental design of reporter assays.

## Resolving the molecular mechanisms of non-coding variants using plasmid Fiber-seq

Having established that plasmids are capable of recapitulating endogenous chromatin architectures of regulatory elements, we next sought to determine whether plasmid Fiber-seq could be used to accurately characterize the molecular mechanisms by which non-coding variants cause disease. We have previously identified a SLC39A4 promoter variant (c.-169A>G) that causes acrodermatitis enteropathica (AE) via disruption of a conserved CCAAT box^29^. We find that when this region is present within a plasmid luciferase reporter, this variant is associated with decreased transcription in both HEK293 and HepG2 cells (**Extended Data** Fig. 11). The WT luciferase activity from the plasmid was significantly higher in HEK293 cells (derived from fetal kidney) than in HepG2 cells (derived from liver), despite the fact that *SLC39A4* encodes a zinc transporter that is most highly expressed in liver tissue. We evaluated if plasmid Fiber-seq could resolve both how the promoter variant decreased transcriptional activity and why HEK293 showed increased expression over HepG2.

Application of plasmid Fiber-seq to HepG2 and HEK293 cell lines co-transfected with the WT and variant luciferase constructs revealed that the pSLC39A4 plasmid contains a large NFR directly overlapping the *SLC39A4* promoter that was unchanged along the plasmids containing the c.-169A>G variant (**Fig. 5a**). A quantitative comparison of protein occupancy differences between the WT and c.-169A>G variant constructs revealed that the variant was associated with a distinct change in the occupancy of 20-70 bp footprints directly overlapping the A>G variant (**Fig. 5b)**. Specifically, a 20-40 bp footprint at this site was abrogated on the variant plasmid in both HepG2 cells and HEK293 cells, consistent with this variant disrupting NF-Y factor occupancy at this CCAAT box. Furthermore, a 40-70 bp footprint at this site was also abrogated on the variant plasmid in HEK293 cells and to a lesser degree HepG2 cells, consistent with this variant disrupting Pol II complexes from assembling at this CCAAT box. Of note, on the WT *SLC39A4* construct this Pol II-sized 40-70 bp footprint showed greater occupancy in HEK293 cells (**Fig. 5b)**, indicating that formation of this Pol II-sized complex may be responsible for the increased transcription of the WT *SLC39A4* construct in HEK293 cells. Overall, these experiments demonstrate that this variant disrupts the ability of an activator TF from occupying the CCAAT box, and that differences in the luciferase activity of this construct in HepG2 and HEK293 cells may reflect the differential ability of CCAAT-binding factors in these cells to recruit and assemble Pol II complexes at this site.

**Figure 5.**
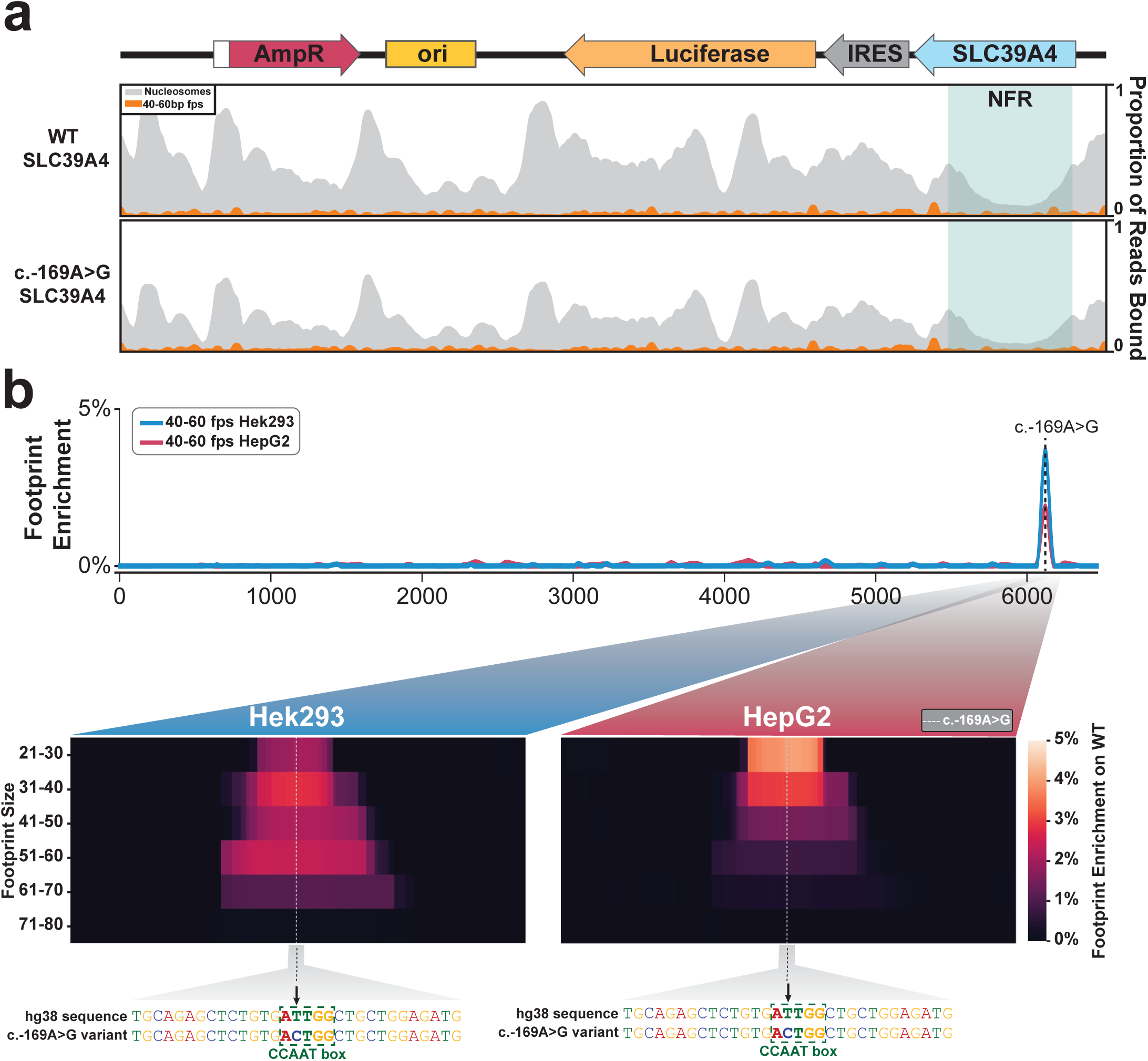
| Fiber-seq of a pathogenic variant in the SLC39A4 promoter. **(a)** Smoothed line plot of pSLC39A4 containing either the Wild-type (top) or Variant (bottom) SLC39A4 promoter sequence in HEK293. Blue box highlights the nucleosome-free region (NFR) present over the wild-type and variant promoter sequence in both HEK293 and HepG2 cell types. Nucleosome occupancy is shown in grey, and 40-60 bp footprints (fps) are shown in orange. **(b)** smoothed line plot showing the location of 40-60 bp footprints enriched on the wild-type promoter sequence relative to the variant sequence in HEK293 (blue) and HepG2 (red) cells.The dashed vertical line indicates the position of the c.-169A>G variant. Heatmaps show the range of footprint sizes that are enriched on the wild-type sequence relative to the variant sequence in HEK293 and HepG2 cells, with footprints binned in 10 bp increments. Heatmaps are centered over the CCAAT box where the c.-169A>G variant is located, highlighted in the underlying sequence.

Having established that plasmid Fiber-seq can help elucidate the molecular mechanisms underlying pathogenic non-coding variants, we next evaluated whether plasmid Fiber-seq could be combined with massively parallel reporter assays (MPRAs) to elucidate variant effects in a high-throughput manner. MPRAs are a widely used plasmid-based approach to study the effect of variants on regulatory element activity, providing insight into variants that increase or decrease transcript abundance. While MPRAs have proven valuable in identifying variant effects, they provide little insight into the molecular mechanisms driving the change in activity (*i.e.,* is the effect mediated by altered occupancy of a transcriptional activator, transcriptional repressor, or altered RNA stability). Importantly, these distinct molecular mechanisms have dramatically different therapeutic implications.

We performed plasmid Fiber-seq on a saturation mutagenesis MPRA library of the low-density lipoprotein receptor (*LDLR*) promoter. Pathogenic variants in the LDLR promoter are known to be causative of familial hypercholesterolemia (FH)^30^. This MPRA library has previously been shown to contain clusters of variants that result in a consistent increase or decrease in transcript abundance^31^ (**Fig. 6a**), but the mechanisms mediating these changes have been largely unknown. We sequenced this plasmid Fiber-seq library using only 60% of a single PacBio Revio SMRT cell, which achieved 1,505,886 full length plasmid reads. As plasmid Fiber-seq is grounded in highly accurate long-read sequencing, it synchronously permits us to identify the variants present on each sequenced construct, as well as their associated overlying chromatin patterns. In total, 950 out of 954 possible SNVs across the 318 bp *LDLR* promoter were present in our sequencing reads (>99% SNV coverage). However, as the MPRA variant library was generated using an error-prone PCR approach, 19% of the reads contained the WT *LDLR* promoter sequence, and only 127 (13.4%) variants had >1000 plasmid fibers containing only that variant, meeting our minimum coverage cutoff for footprint analysis. To identify protein occupancy patterns associated with altered transcript abundance we combined the MPRA transcript results with the plasmid Fiber-seq footprints. In the first example, the c.-223A>G variant causes a 2-fold decrease in transcript abundance, and plasmid Fiber-seq showed that this variant causes the loss of a 20-60 bp footprint that selectively occupies this location on the WT plasmid (**Fig. 6b**). Based on these findings, it appears that this variant causes decreased *LDLR* expression by disrupting the occupancy of a transcriptional activator. In contrast, the c.-214T>C variant causes a 1.57-fold increase in transcript abundance, and plasmid Fiber-seq showed that this variant causes the gain of a 20-60 bp footprint that selectively occupies this location on the variant plasmid (**Fig. 6b**). Based on these findings, it appears that this variant causes increased *LDLR* expression by increasing the occupancy of a transcriptional activator. Of note, this effect is highly sequence specific, as the c.-214T>A variant shows no change in footprint abundance at this site, and actually corresponds to a 0.89-fold decrease in transcript abundance (**Extended Data** Fig. 12).

**Figure 6.**
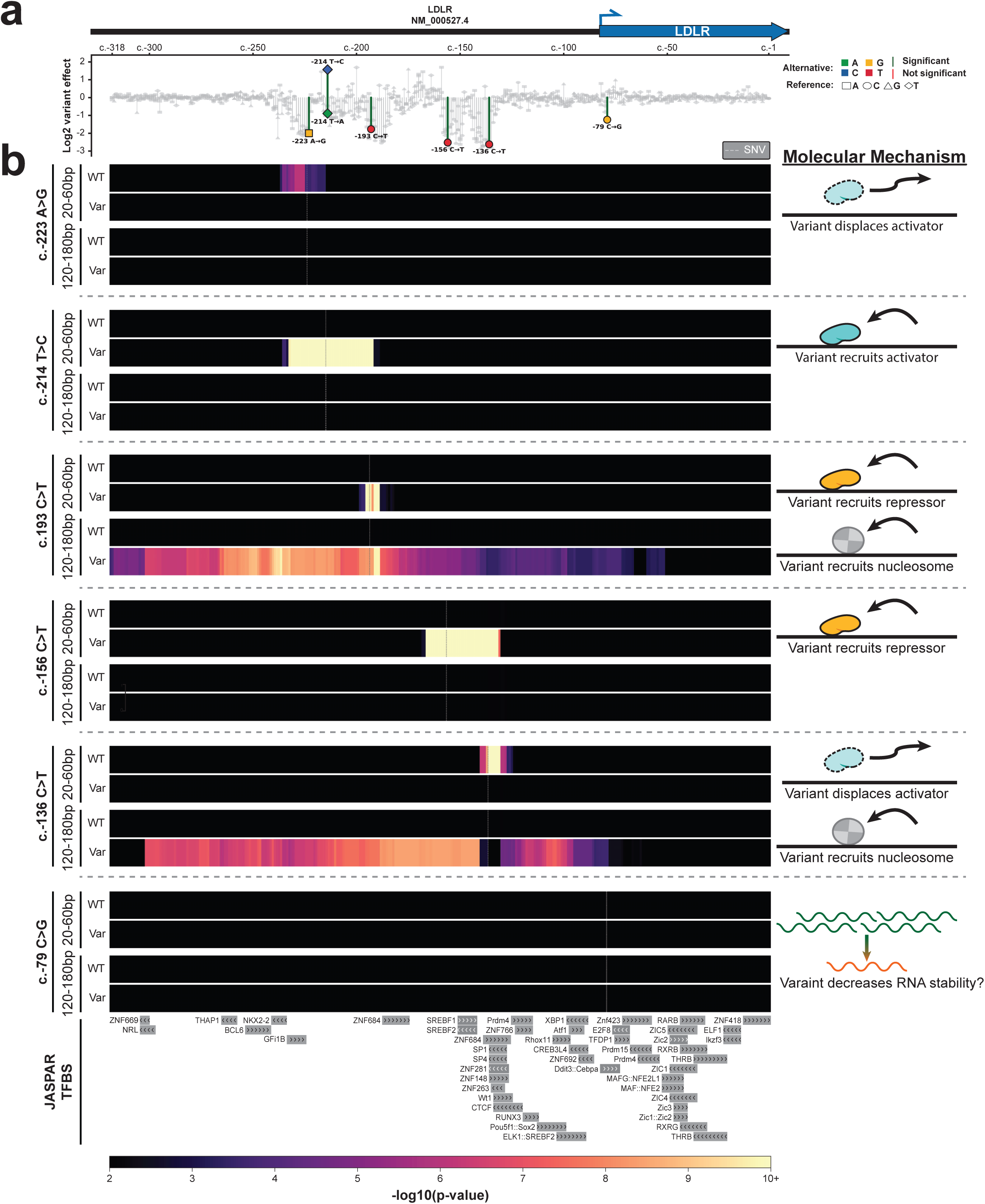
| Molecular mechanisms of variant effects. **(a)** Log2 variant effects of all SNVs ordered by their RefSeq transcript position in NM_000527.4 of the hypercholesterolemia-associated LDLR promoter^31^. Colored data points correspond to variants whose molecular mechanism of variant effect is defined in b. Significance level (red/green lines) is 10^-5^. **(b)** Heatmaps showing the enrichment of protein footprint occupancy for both small proteins (20-60p) and nucleosomes (120-180 bp) along the LDLR promoter for both the wild type and specified variant sequence for six individual variants. Color intensity corresponds to the -log10(p-value) of the enrichment, with a lower cutoff corresponding to a false discovery rate of < 5%. Cartoon to the right of each variant diagrams the molecular mechanism by which the variant alters promoter activity. JASPAR^40^ transcription factor binding motifs (TFBS) along the bottom highlight the abundance of transcription factor binding motifs within this promoter region.

In addition to resolving occupancy patterns of transcriptional activators, plasmid Fiber-seq also exposed variants whose negative effects on expression appear to be mediated by transcriptional repressors. For example, the clinical variant c.-156C>T^32^ causes a 2.52-fold decrease in transcript abundance, and our plasmid Fiber-seq data showed that this variant causes the gain of a 20-60 bp footprint that selectively occupies this location on the variant plasmid (**Fig. 6b**). Based on these findings, it appears that this pathogenic variant causes decreased *LDLR* expression by increasing the occupancy of a transcriptional repressor at this site. We find additional evidence of this phenomena with the c.-193C>T variant. This variant causes a 1.77-fold decrease in transcript abundance, and our plasmid Fiber-seq data showed that this variant causes the gain of a 20-60 bp footprint along with a 120-180 bp footprint that selectively occupy this location on the variant plasmid (**Fig. 6b**). Based on these findings, it appears that this variant causes decreased *LDLR* expression by increasing the occupancy of a transcriptional repressor and a nucleosome at this site.

Lastly, plasmid Fiber-seq also exposed variants whose effects do not appear to be mediated by gene regulatory changes. Specifically the c.-79C>G variant causes a 1.24-fold decrease in transcript abundance, and our plasmid Fiber-seq data showed that this variant causes no change in footprint occupancy along the plasmid (**Fig. 6b**). Notably, this variant is located immediately downstream of the *LDLR* transcription start site (TSS), and based on these findings, it appears that this variant likely causes decreased *LDLR* expression by decreasing *LDLR* RNA stability. Overall, these findings demonstrate that plasmid Fiber-seq can resolve the molecular mechanisms of non-coding variants, thereby expanding the utility of plasmid-based reporter assays.

## Discussion

In summary, plasmids are indeed capable of recapitulating nuclear genome-encoded chromatin architectures, and the focal occupancy of nucleosomes and transcription factors along transfected plasmids is central to their transcriptional activity within mammalian cells. However, plasmids consistently adopt a heterogeneous and incomplete chromatin architecture relative to nuclear-encoded chromatin fibers, and surrounding sequence context plays an integral role in modulating whether a plasmid-encoded regulatory element will recapitulate nuclear genome-encoded chromatin architectures.

Importantly, we establish that plasmid Fiber-seq can comprehensively measure the chromatin architecture of full-length transfected plasmid molecules with near base-pair resolution, thereby illuminating whether a regulatory element adopts a distinct chromatin architecture when present within the plasmid or nuclear genome context. Plasmid Fiber-seq accommodates a wide range of full-length plasmid construct sizes and compositions while synchronously capturing nuclear-encoded chromatin fibers that serve as a robust internal control. The high read coverage (millions of individual plasmid reads) obtained from a single experiment enables two significant advantages: first, the depth of coverage permits the high-resolution detection and quantification of DNA-occupying proteins, ranging from TFs to larger protein complexes and nucleosomes. Second, the high-throughput capacity allows for the simultaneous analysis of thousands of plasmid constructs in a single reaction, making this approach compatible with large-scale functional genomic assays. However, our findings do raise concerns regarding the current standard of using only short (*i.e.,* ∼300 bp) elements in these large-scale functional genomic assays. Specifically, we find multiple examples where additional genomic sequence context would have enabled a short element to accurately recapitulate the genomic chromatin architecture of that element. It remains to be seen how much this feature contributes to the high false negative rate of MPRA screens of genetic variants in disease-associated loci^33,34^. However, it is also notable that for some elements, additional genomic sequence context negatively impacted the ability of the plasmid to accurately recapitulate the genomic chromatin architecture of that element, highlighting the role of plasmid Fiber-seq for validating plasmid-based reporter assays.

We find that transcription-associated footprints (*i.e.,* TF and Pol II sized footprints) within plasmid-encoded regulatory elements are present along both lowly chromatinized and highly chromatinized plasmids, and that the abundance of these footprints varies based on a plasmid’s overall chromatin state. This finding suggests that the majority of the transcript signal from bulk plasmid-based reporter assays, even at the single-cell level^13^, represent an average of transcriptional activity across diverse plasmid chromatin states rather than a functional readout of a uniformly chromatinized plasmid state. Furthermore, as the lowly chromatinized state, by definition, is not adopting a chromatin architecture that recapitulates nuclear genome-encoded chromatin, this will cause plasmid-based assays to stray in their estimation of how a regulatory element would function in its endogenous location. In contrast to traditional transcript readouts of plasmid-based assays, plasmid Fiber-seq can illuminate the protein-DNA occupancy mechanisms underlying variant effects, including resolving the occupancy of transcriptional activators and repressors within regulatory elements. This has the potential to improve how we therapeutically target genetic diseases. For example, transcriptional repressors have proven to be fruitful therapeutic targets (*e.g.,* BCL11A in the treatment of hemoglobinopathies^35^), and using plasmid Fiber-seq we have identified putative repressors that may be regulating the *LDLR* locus. Heterozygous loss-of-function pathogenic variants in *LDLR* cause FH, and our data suggests that targeting these putative *LDLR* repressors may be a fruitful therapeutic strategy for FH.

The heterogeneous chromatinization we observe likely reflects the steady-state chromatin pattern of plasmids within mammalian nuclei. Although it is possible that some of the completely unchromatinized plasmid molecules may derive from plasmids wrapped by a nuclear envelope like structure that is being retained during the nuclei isolation step^24^, the far majority of lowly chromatinized plasmids contain well-positioned nucleosome and sub-nucleosome-sized footprints in a manner that indicates that they are actively interacting with the nucleoplasm, and are therefore likely of nuclear origin. Consequently, the presence of these lowly chromatinized fibers indicates that plasmids generally adopt a distinct chromatin architecture relative to that of the nuclear genome, as replicated nuclear genome-encoded DNA is often chromatinized in a matter of minutes^36^. It remains to be seen if other sources of exogenous DNA similarly adopt a heterogeneous and incomplete chromatin architecture. For example, many Food and Drug Administration (FDA)-approved gene therapies use adeno-associated viruses (AAV) which are single-stranded DNA viruses that are converted into double-stranded DNA within the nucleus. We anticipate that plasmid Fiber-seq could similarly be adopted to resolve how these AAV vectors are chromatinized, which may provide valuable insights into improving their therapeutic potential.

Overall, plasmid Fiber-seq can elucidate key relationships between sequence, chromatin architecture, and function, thereby guiding how exogenous DNA-based technologies are optimized and expanding the utility of plasmids as a foundational molecular biology tool.

## Acknowledgements

We thank B. Martin and J. Shendure for providing the LDLR saturation mutagenesis library, K. Munson and the University of Washington PacBio Sequencing Services, the Northwest Genomics Center, the University of Washington Laboratory Medicine and Pathology Flow Cytometry Core Facility, C. Parrott and G. Mikol for assistance in plasmid construct assembly, and K. Straus, C. Kwong, A. Hosokai, and N.T. Smith for assisting with plasmid purification.

Funding: A.B.S. holds a Career Award for Medical Scientists from the Burroughs Wellcome Fund and is a Pew Biomedical Scholar. This research is supported by the National Institutes of Health (NIH) grant 1DP5OD029630 to A.B.S. and a Brotman Baty Institute for Precision Medicine Catalytic grant to A.B.S. B.J.M. and L.M.S. were supported by the Center for Multiplexed Assessment of Phenotype, an NHGRI Center of Excellence in Genome Sciences 2RM1HG010461.

## Methods

### Plasmids

pUC19 was a gift from Joachim Messing (Addgene plasmid # 50005; http://n2t.net/addgene:50005; RRID:Addgene_50005)^37^, pEF-GFP was a gift from Connie Cepko (Addgene plasmid # 11154; http://n2t.net/addgene:11154; RRID:Addgene_11154)^20^, and ColorfulCell was a gift from Pierre Neveu (Addgene plasmid # 62449; http://n2t.net/addgene:62449; RRID:Addgene_62449)^38^ . The LDLR saturation mutagenesis library was a gift from B. Martin from the Shendure lab^31^. Wild-type and variant *SLC39A4* luciferase constructs were generated as previously described^29^.

Deletion variants of pEF-GFP (Partial-ΔEF1α, Full-ΔEF1α, and Full-ΔSV40) were generated via PCR amplification with opposite-facing primers flanking deletion regions. The barcoded pEF-GFP series (G52G, R97R, E143E, D181D) was created through site-directed mutagenesis with primers containing synonymous substitutions within the GFP gene (G52G, R97R, E143E, D181D). All constructs were generated via the Q5® Site-Directed Mutagenesis kit (NEB E0554) and transformed into DH5α.

The pIRES-GFP backbone was generated by replacing the EF1α promoter in pEF-GFP with a 755 bp gBlock containing a multiple cloning site, splice acceptor sequence, stop codons in all three reading frames, and an IRES element using Gibson assembly (NEB E2611).

The pDPAGT1-GFP construct was created by amplifying the DPAGT1 promoter from HEK293 genomic DNA using primers with 40 bp overlapping homology arms, followed by Gibson assembly with the linearized pIRES-GFP backbone.

Additional promoter-GFP constructs (pWASF1, pUBA1, pMYC, pNAPA, pFLCN, pHBB, pMYBPC3, and pMYH7) were generated by amplifying respective promoter sequences from HEK293 genomic DNA with restriction site-containing primers. WASF1, UBA1, MYC, and MYH7 were cloned into the SalI/EcoRI sites, while NAPA, FLCN, HBB, and MYBPC3 were cloned into the XbaI/SalI sites. Following digestion and purification, promoter fragments were ligated into the pIRES-GFP vector using T4 ligase (NEB M2200).

Constructs containing 300 bp promoter segments were generated by inserting synthetic 350 bp gBlocks (300 bp core sequence with flanking homology arms, IDT) into SalI/XbaI-digested pIRES-GFP using HiFi DNA Assembly (NEB E5520). Specific primers and genomic coordinates for cloned regions are provided in Extended Data table 1.

## Bacterial strains and culture conditions

All bacterial strains used in this study were grown in Lysogeny Broth (LB) at 37 °C or on LB medium solidified with agar (RPI, cat# L24030-100.0). Filter sterilized ampicillin (ThermoFisher Cat # J60977.14) (100 mg/L for plasmid propagation), was added to culture when necessary. E. coli strains DH5α (NEB, cat# C2987H), *dam–/dcm–* (NEB, cat# c2925H), and ER2796 (gift from NEB) were used to propagate plasmid DNA.

## Transformation and extraction of plasmid constructs from ER2796

The methyltransferase deficient E. coli strain ER2796 (NEB) was made chemically competent using the Mix & Go! E. coli Transformation Kit (Zymo Research T3001), and transformations were performed according to the manufacturer’s instructions. Plasmid preparations were performed via mini-prep (NEB T1010) or maxi-prep (Zymo Research D4202) from overnight cultures grown in 5 mL or 150 mL LB media + 100 ug/mL ampicillin (ThermoFisher 60977.14).

## Cell Cultures

All cell lines were maintained at 37°C with 5% CO₂ and passaged at 80-90% confluency using 0.25% trypsin-EDTA (Fisher, 25200056). HEK293, HepG2, and A375 cells were cultured in DMEM (Fisher 11995040) with 10% FBS (FBS, Fisher 10082147) and 1% penicillin-streptomycin(Fisher 15070063), while HAP1 cells were maintained in IMDM (Fisher 12440046) with the same supplements. Cells were cryopreserved at 4-5×10⁶ cells/mL in FBS with 10% DMSO (Sigma Aldrich D2650) and stored in liquid nitrogen. For thawing, cryovials were rapidly warmed in a 37°C water bath and cells were pelleted, resuspended in fresh medium, and transferred to culture vessels.

## Generation of relaxed pEF-GFP

pEF-GFP sourced from ER2796 was digested with Nt.BspQI (NEB R0644) and re-ligated by T4 ligase (M2200) according to the manufacturer’s protocol. Ligated DNA was purified using a DNA cleanup kit (NEB T1130).

## Flow cytometry of allelic series

Flow cytometric analysis was performed on a BD FACSymphony A3 Cell Analyzer (BD Biosciences) using the 488 nm laser with a 530/30 nm bandpass filter for GFP detection. For each sample, 10,000 events were collected using BD FACSDiva software. Data were analyzed with FlowJo software (BD Biosciences), employing a sequential gating strategy to identify single, viable, GFP-positive cells. GFP expression was quantified by the percentage of positive cells with a fluorescence intensity above that of non-transfected HEK293 cells.

## Plasmid Fiber-seq Reactions

Nonspecific methyltransferase, Hia5, was purified, and the activity was quantified as previously described^14^. For each experiment, cells were seeded at 2-3×10⁶ cells per 10 cm plate on the day prior to transfection. Cells were transfected using Lipofectamine LTX (ThermoFisher #15338100) following the manufacturer’s protocol. In brief, 10 μg plasmid DNA was diluted in 1 mL Opti-MEM medium (Fisher 31985070) and supplemented with 10 μL PLUS Reagent (1:1 ratio to DNA). The mixture was gently mixed and incubated at room temperature for 10 minutes. Subsequently, 30 μL Lipofectamine LTX reagent (3:1 ratio to DNA) was added, and the transfection complex was incubated at room temperature for 30 minutes. The transfection complex was added dropwise to each plate and mixed by gentle shaking. Cells were incubated with the transfection reagent until harvest.

Harvested cells were resuspended in Buffer A (15 mM Tris-Cl, pH 8, 15 mM NaCl, 60 mM KCl, 1 mM EDTA pH 8, 0.5 mM EGTA pH 8, 0.5 mM spermadine) and aliquoted at 2×10⁶ cells per 60 μl. Cell lysis was performed by adding 60 μL of lysis buffer (Buffer A supplemented with 0.05% IGEPAL), mixed by flicking, and incubating on ice for 10 min. Nuclei were pelleted at 350×g for 5 min at 4°C and resuspended in 60 μL Buffer A supplemented with 0.8 mM SAM and 100 U of Hia5 methyltransferase. After incubation at 25°C for 10 min, the reaction was stopped by adding SDS to a final concentration of 1%. Plasmid DNA was extracted by adding 137 μL Monarch Buffer B1 (NEB T1111) and purified by mini-prep (NEB T1110) according to the manufacturer’s protocol, with elution in 20 μL molecular-grade water. Purified plasmid DNA was linearized by adding 20 U restriction enzyme, 5 μL 10× rCutSmart Buffer (NEB), and ddH₂O to 50 μL, followed by incubation at 37°C for 2 h. Linearized DNA was purified using the Monarch Spin PCR & DNA Cleanup Kit (NEB) and eluted in 20 μl.

All plasmid Fiber-seq reactions were performed in triplicate performing separate transfections on separate days, with the exception of the 5mCpG methylated pEF-GFP, which was performed in duplicate.

## PacBio Library Preparation and Sequencing

Purified DNA samples were quantified using the Qubit dsDNA high-sensitivity assay (Qubit Q32851) following the manufacturer’s protocol. Multiplexed library preparation was performed using the SMRTbell prep kit 3.0 (PacBio, #102-141-700) and SMRTbell barcoded adapter plate 3.0 (PacBio, #102-009-200) according to the manufacturer’s protocol. Prepared libraries were loaded onto a single Revio SMRT cell (R1 chemistry) and sequenced by the University of Washington PacBio Sequencing Services.

## Fiber-seq Data Processing

m6A methylation and nucleosome footprints were identified using fibertools-rs^18^, and CpG methylation was called using jasmine (PacBio). For genomic samples, Fiber-seq inferred regulatory elements (FIRE) were called using the FIRE pipeline^39^. For plasmid samples, sub-nucleosomal footprints were called using FiberHMM^21^. Plasmid reference sequences were obtained from whole plasmid sequencing performed by Plasmidsaurus using Oxford Nanopore Technology with custom analysis and annotation, and plasmid Fiber-seq reads were aligned to a reference containing both linearized plasmid reference sequences and the hg38 reference genome. For downstream plasmid analysis, reads were filtered for full-length fibers (reference length ±50 bp) with ≥10% m6A methylation (**Extended Data** Fig.13).

## Comparison of plasmid and genomic nucleosome density

pEF-GFP plasmid fibers and genomic fibers were sourced from the same plasmid Fiber-seq reaction. Genomic reads were filtered for fibers with identical length (+/- 50 bp) to the linearized pEF-GFP plasmid, and were required to contain at least two FIRE peaks within the read to reflect the accessibility conferred by the EF1α promoter on pEF-GFP. Nucleosome footprints for both read sets were called using fibertools-rs^18^

## Categorization of lowly, intermediately, and highly chromatinized plasmid reads

Plasmid chromatin states were classified using a benchmark slope of 0.005 nucleosomes/bp derived from plotting nucleosome density against genomic fiber read length. Reads with <20% of the theoretical nucleosome density were classified as lowly chromatinized, >60% as highly chromatinized, and the remainder as intermediately chromatinized.

## Encode tracks for CTCF and DNaseI footprints for genomic promoters

The following ENCODE (ENCODE Project Consortium 2012) tracks were included: CTCF ChIP-seq of HEK293, ENCSR000DTW; DNase-seq of HEK293T ENCSR000EJR (**Fig. 4**).

## Plasmid Fiber-seq of the LDLR MPRA Library

The LDLR library was bottlenecked 1:1000 before being transformed into the ER2796 bacterial cell line via electroporation. The full contents of the electroporation were added to 150mL overnight culture of LB supplemented with 100 ug/mL ampicillin, and extracted via maxi-prep (Zymo Research D4202). The LDLR MPRA library was transfected into HEK293 cells (**Extended Data** Fig. 14), with plasmid Fiber-seq being performed 48h post-transfection.

## LDLR MPRA Statistical analysis

We used FiberHMM to detect protein footprints along both the WT and variant reads. The large number of WT reads allowed us to generate statistical comparisons between WT and variant footprint occupancy.

To establish a control dataset, we generated a WT dataframe containing the average proportion of reads with a given footprint size along each base of the promoter region, placing the footprint sizes in 10 bp bins. We subsampled 5,000 random reads from our pool of 291,888 WT reads 10,000 times. For each subsample, we calculated the average proportion dataframe, with our final dataframe representing the average value of our 10,000 subsamples.

For each variant with at least 1000× coverage, we generated an equivalent dataframe, calculating the proportion of reads with specific sized footprints at each position along the LDLR promoter. We then calculated the difference between the WT and variant dataframes in both directions: WT-variant (enrichment of footprints on the WT sequence relative to the variant) and variant-WT (enrichment of footprints on the variant sequence relative to the WT).

To determine statistical significance, we created a null distribution of proportional differences from our WT reads. We again subsampled our WT reads, taking two subsamples of 5,000 reads each, calculating the occupancy proportion dataframes for each, and subtracting them to get the difference in footprint occupancy within our WT samples. This process was repeated 10,000 times to generate a null distribution for each binned footprint size at each position along the promoter region.

For each variant-WT comparison, we calculated a z-score for the observed proportional difference using the mean and standard deviation of the corresponding null distribution for each footprint bin. We then performed a one-tailed p-value test to generate a p-value for footprint enrichment on either the WT or variant sequence. Finally, we applied a Benjamini-Hochberg multiple testing correction with a false discovery rate threshold of 5%. The -log10 of the p-value was used to generate the final enrichment heatmaps.

## Code Availability

The code to reproduce figures and analyses in this study is available on GitHub: https://github.com/StergachisLab/Plasmid-Fiber-seq

---

**Extended Data Figure 1.**
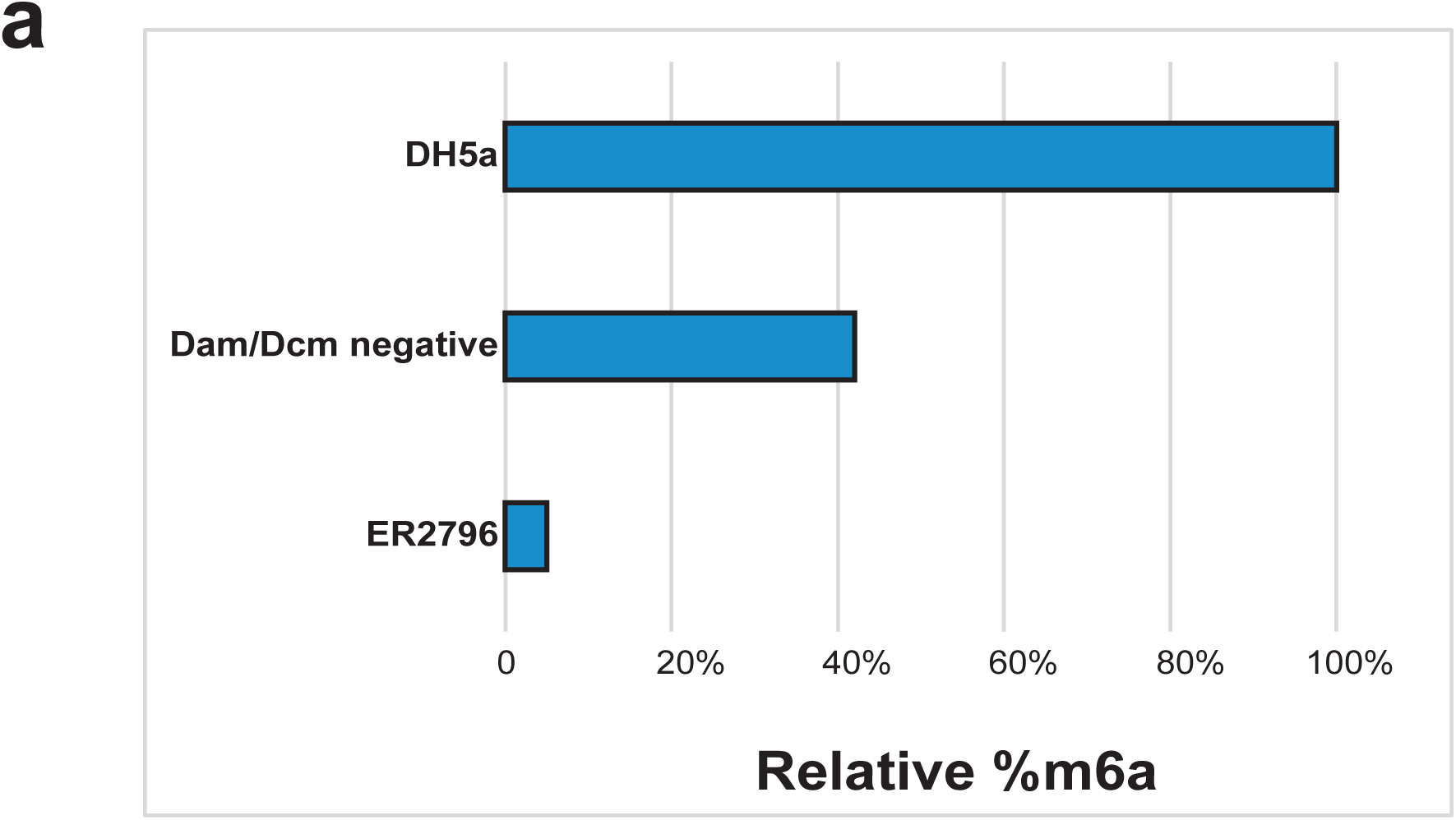
| Removal of m6a from bacterial host **(a)** Quantification of plasmid m6a methylation derived from three different bacterial host strains normalized to the methyltransferase sufficient DH5α strain as measured by small molecule mass spectrometry, performed as previously reported^18^.

**Extended Data Figure 2.**
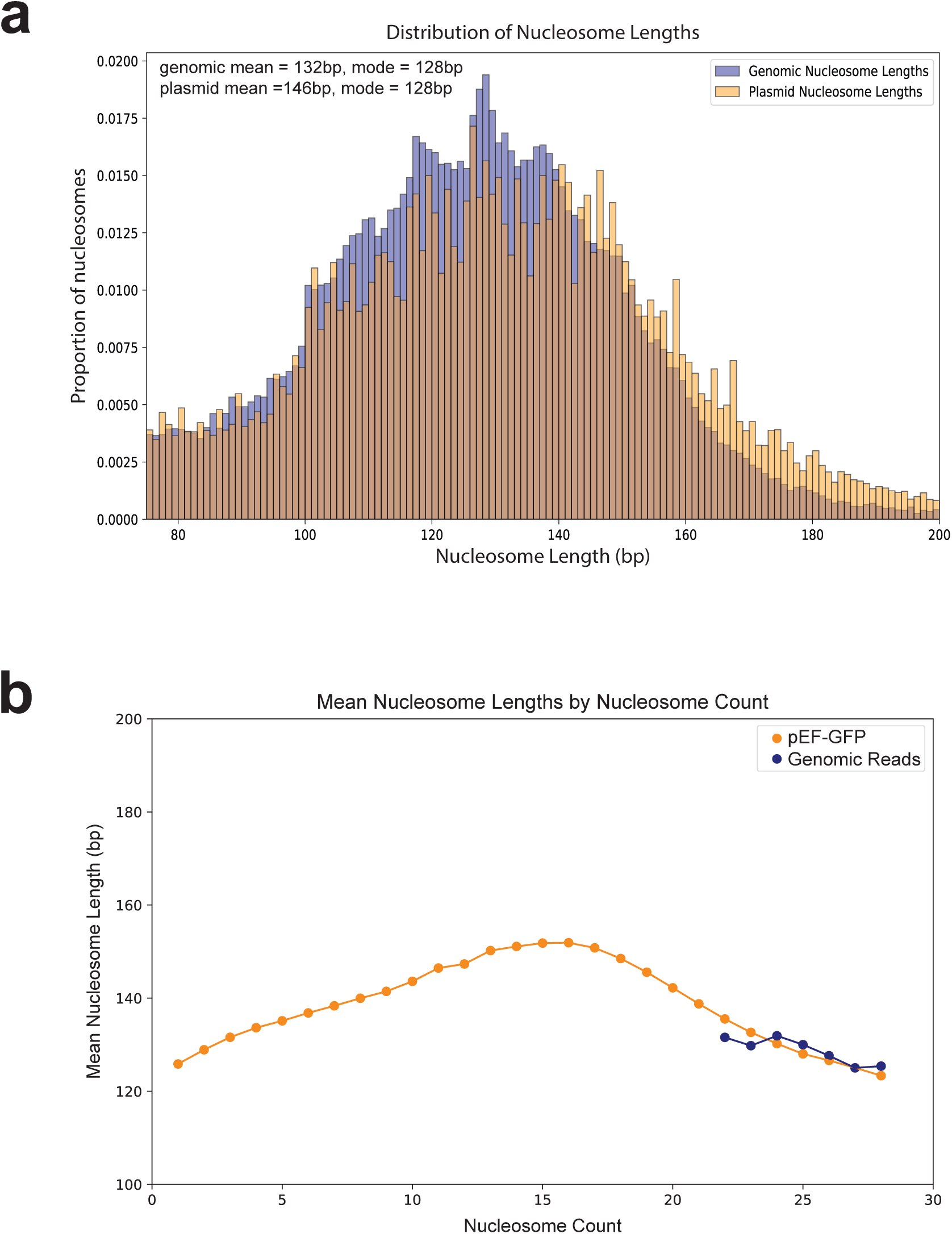
| Comparison of nucleosome sizes between plasmid and genomic fibers at different chromatin states **(a)** A histogram showing the distribution of nucleosome sizes for 87,179 nucleosomes from plasmid and genomic reads sourced from the same Fiber-seq reaction. **(b)** A line plot of the mean nucleosome size along plasmid and genomic fibers across chromatin states.

**Extended Data Figure 3.**
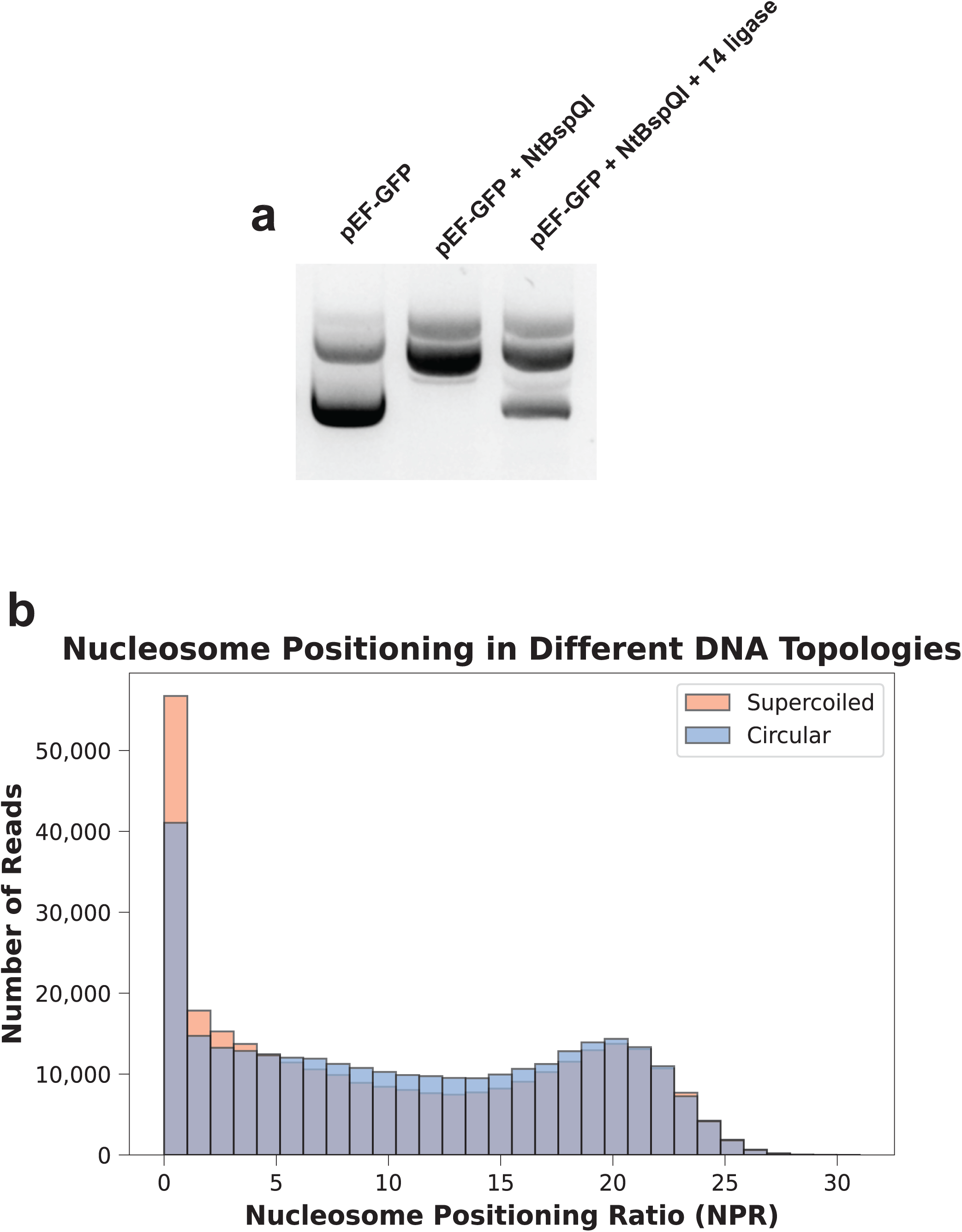
| Supercoiling has minimal effect on plasmid chromatin states **(a)** Agarose gel showing the difference in mobility between supercoiled, nicked, and nicked and re-ligated pEF-GFP, verifying the relaxed state of pEF-GFP. **(b)** Histogram showing the minimal difference in heterogeneous chromatin distribution between supercoiled and circular pEF-GFP 48h post-transfection in HEK293 cells.

**Extended Data Figure 4.**
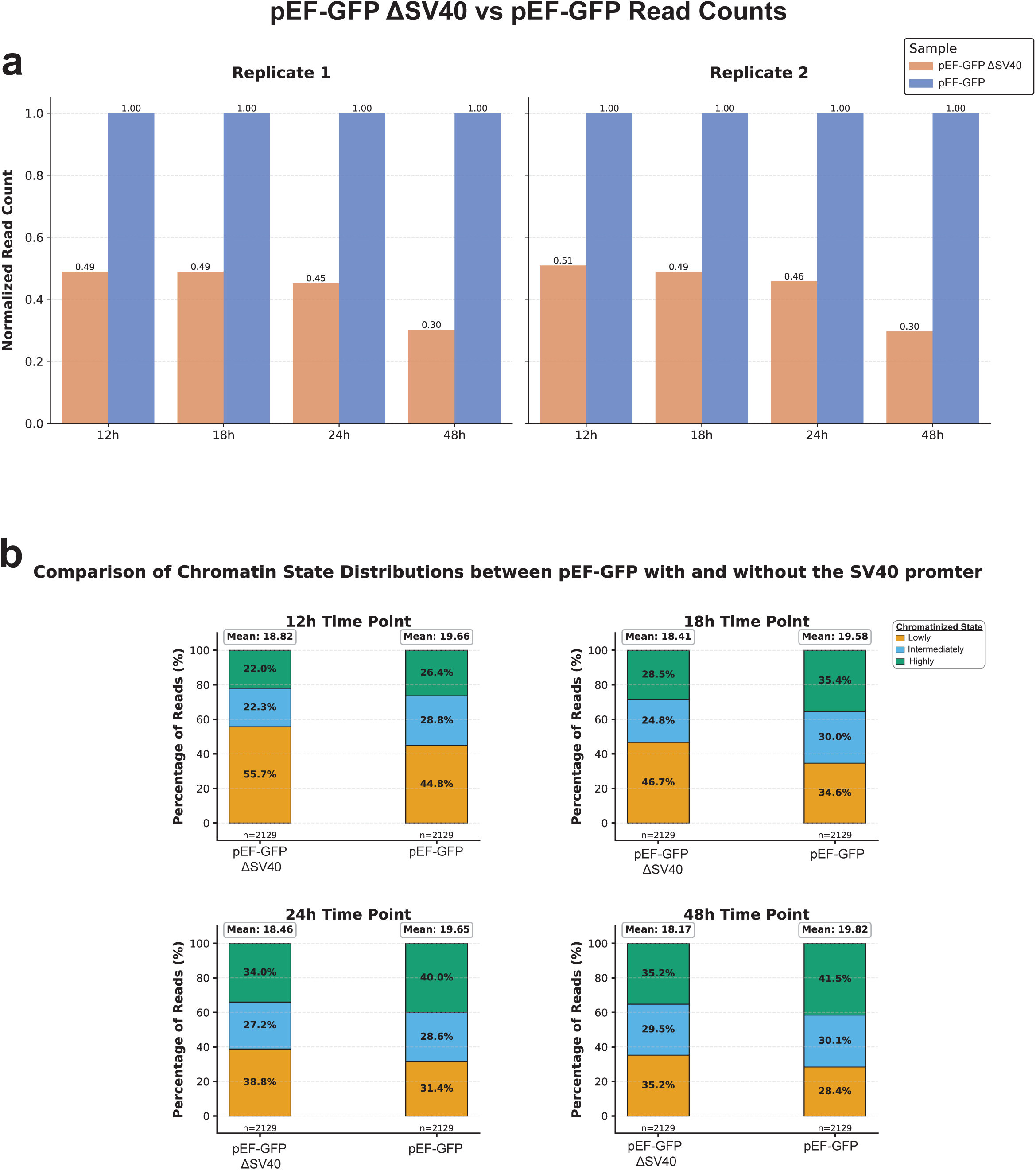
| The SV40 promoter drives plasmid nuclear localization and chromatinization **(a)** Bar plot showing the difference in read counts mapping to pEF-GFP constructs with the SV40 promoter (blue) or without the SV40 promoter (red) 12-, 18-, 24-, and 48-hours after an equimolar transfection in HEK293 cells, normalized to the read count obtained from pEF-GFP containing the SV40 promoter. **(b)** Stacked bar charts comparing the quantity of fibers in the lowly (yellow), intermediately (blue), or highly (green) chromatinized state between the pEF-GFP constructs with or without the SV40 promoter 12-, 18-, 24-, and 48-hours after an equimolar transfection in HEK293 cells. Mean value corresponds to the mean of the highly chromatinized population of fibers.

**Extended Data Figure 5.**
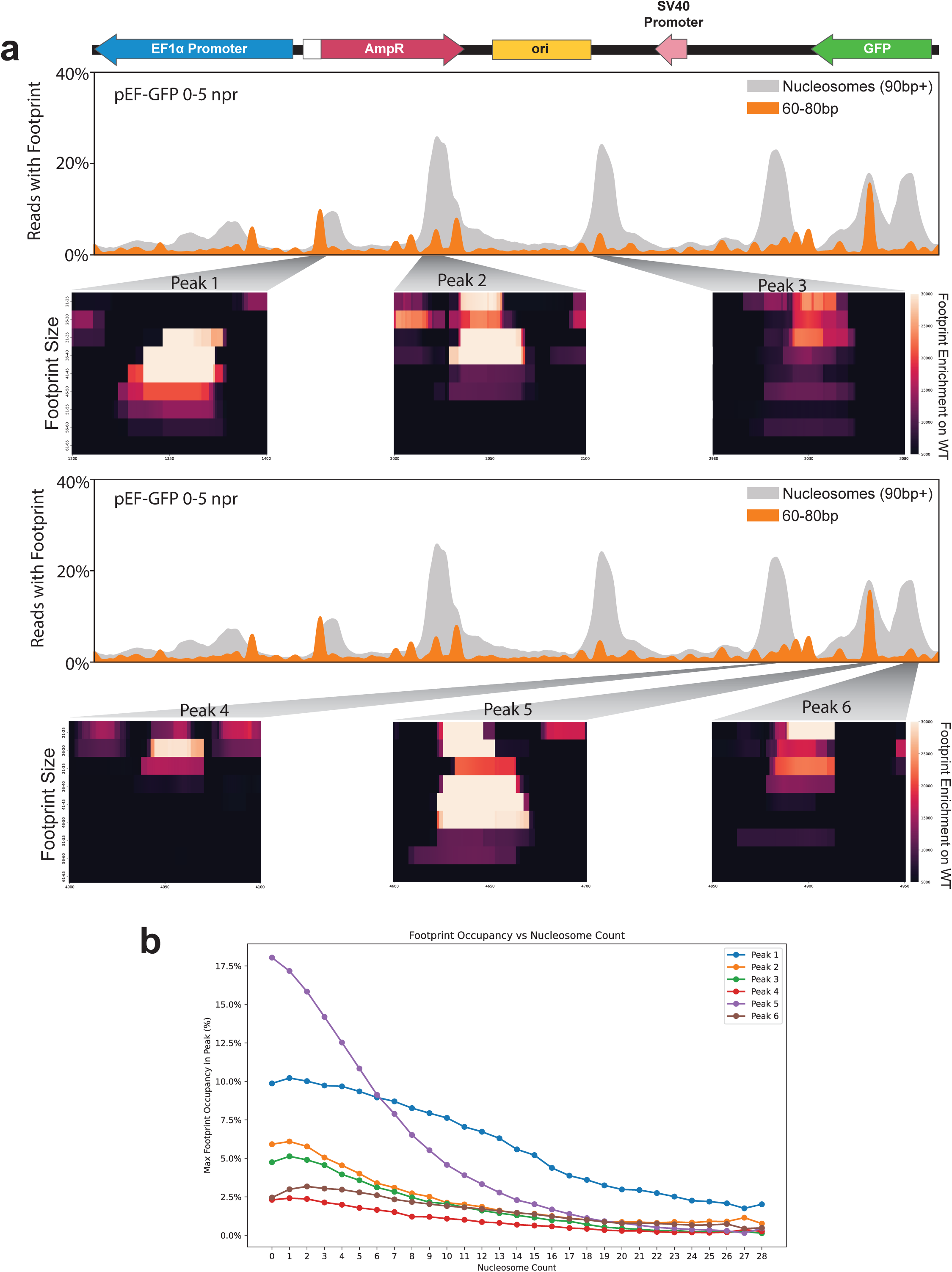
| Nucleating footprints correlate with well-positioned nucleosomes **(a)** Smoothed line plot showing the occupancy of nucleosomes (grey) and 60-80 bp footprints (orange) along the linearized pEF-GFP plasmid. Heatmaps show the diversity of footprint sizes existing along the sequence corresponding to the 40-60 bp footprints peaks that give rise to well-positioned high-affinity nucleosomes, with footprints binned in 5 bp increments. Top and bottom line plots are identical, but separated out to visualize additional heatmaps corresponding to nucleosome nucleating peaks. **(b)** Line plot showing that these small nucleating footprints decrease with increasing nucleosome density.

**Extended Data Figure 6.**
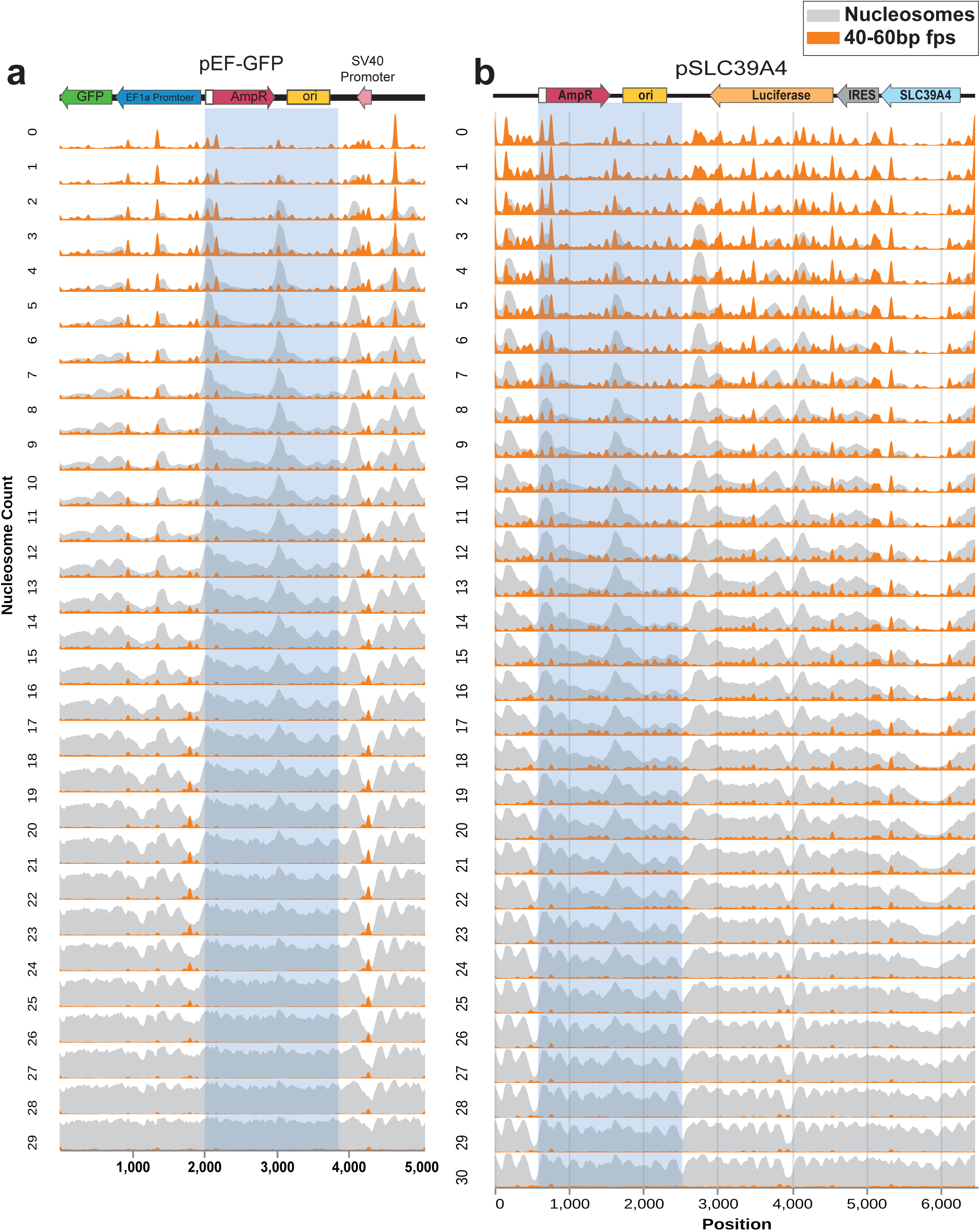
| Nucleosome positioning is sequence specific ridgeplots showing the chromatin architecture of **(a)** pEF-GFP and **(b)** pSLC39A4 across different chromatin states. Blue boxes highlight the shared 1,937 bp prokaryotic backbone, revealing that the small protein nucleation sites and corresponding nucleosome occupancy is highly similar between these two sequences despite differences in their surrounding sequence context.

**Extended Data Figure 7.**
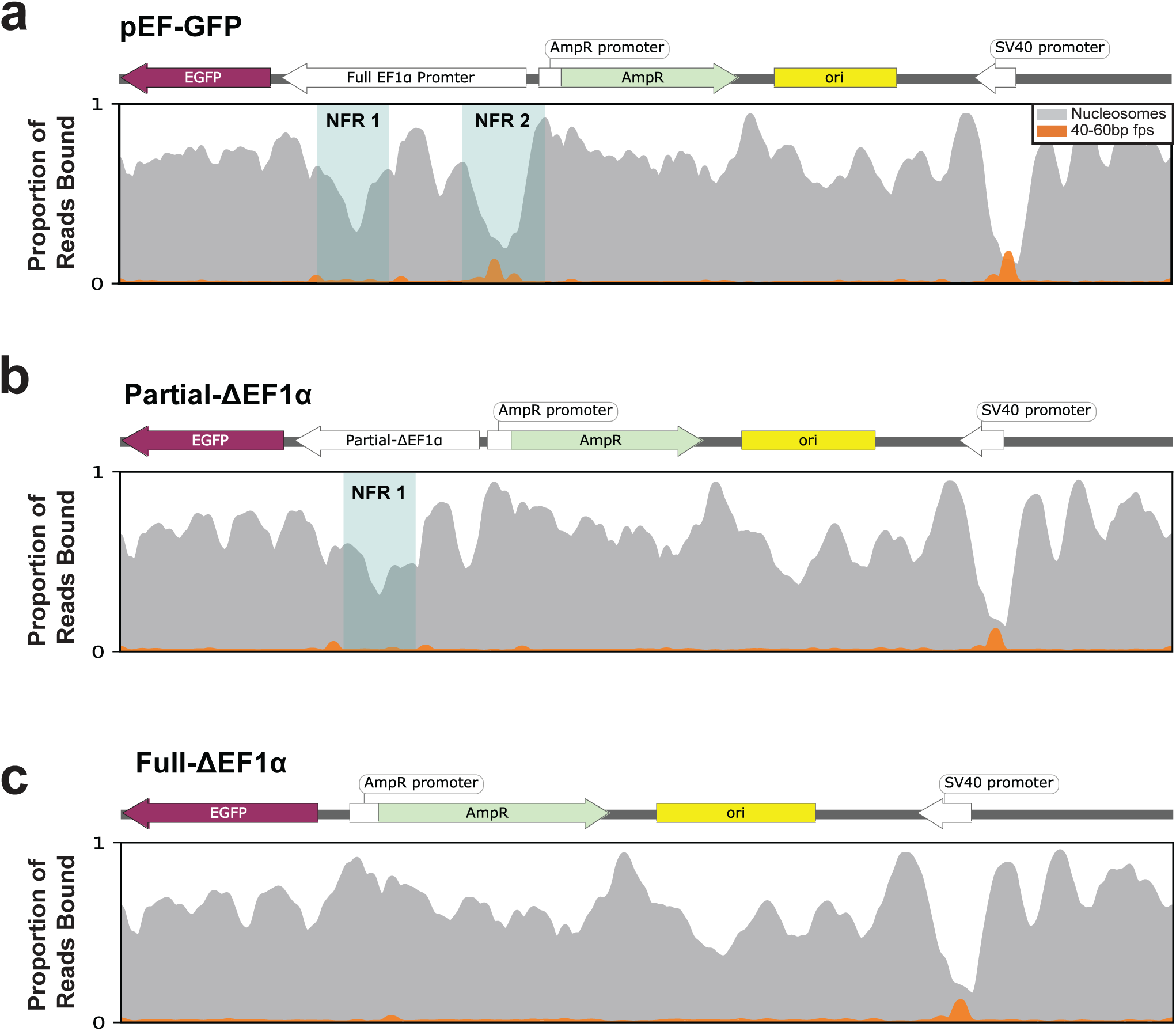
| The partial deletion of EF1α maintains an NFR Smoothed line plot showing the chromatin architecture of **(a)** pEF-GFP, **(b)** Partial-ΔEF1α, and **(c)** Full-ΔEF1α, with the blue box highlighting the nucleosome-free regions (NFR) present on each construct, highlighting the maintenance of NFR1 on the Partial-ΔEF1α construct.

**Extended Data Figure 8.**
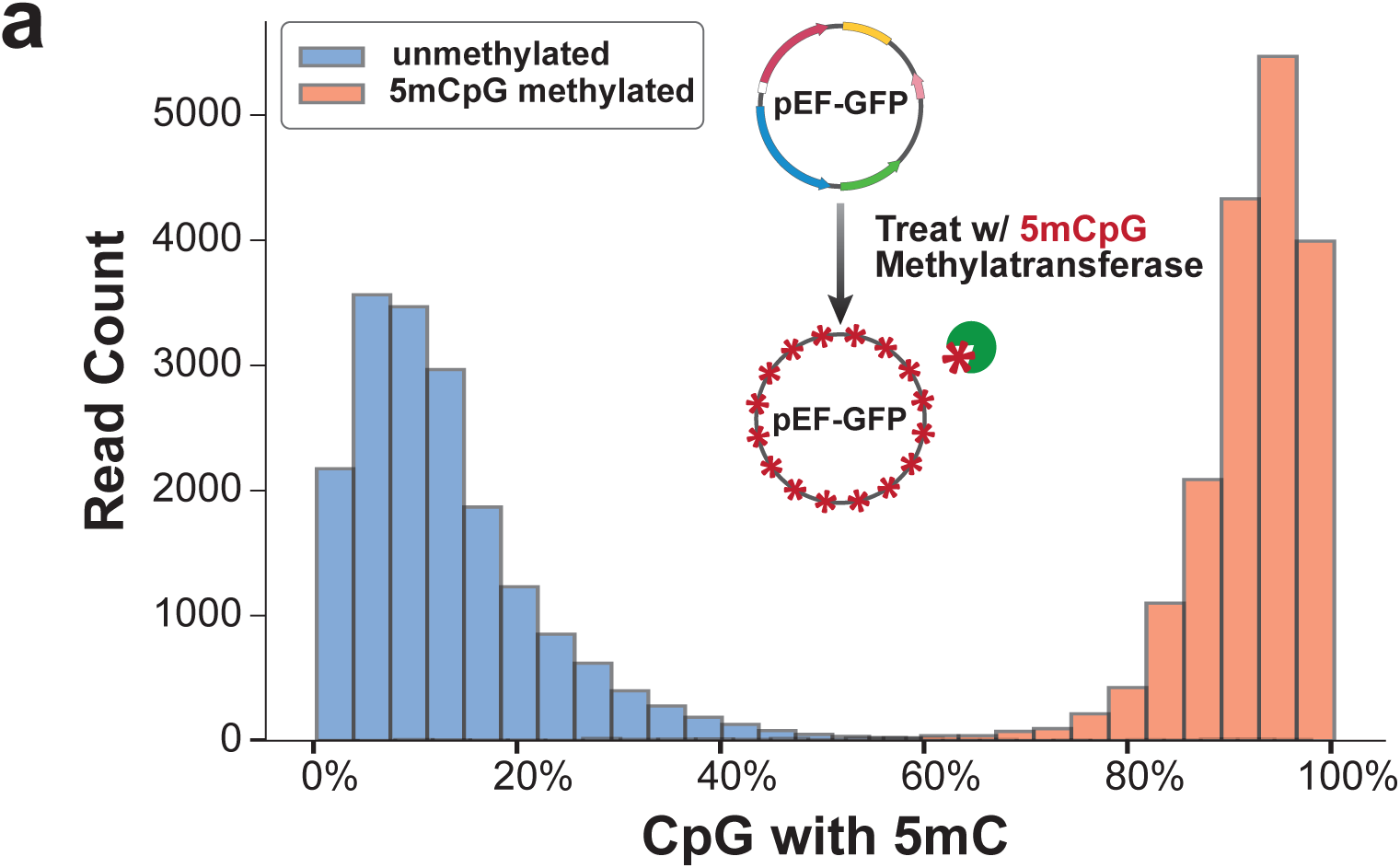
| 5mCpG Methylation of pEF-GFP **(a)** Histogram showing the quantification of 5mCpG methylated CpG dinucleotides for the unmethylated (blue) or 5mCpG methylated (red) pEF-GFP plasmid 48h post-transfection in HEK293 cells. Diagram displays the in-vitro treatment of pEF-GFP with the M.SssI CpG methyltransferase.

**Extended Data Figure 9.**
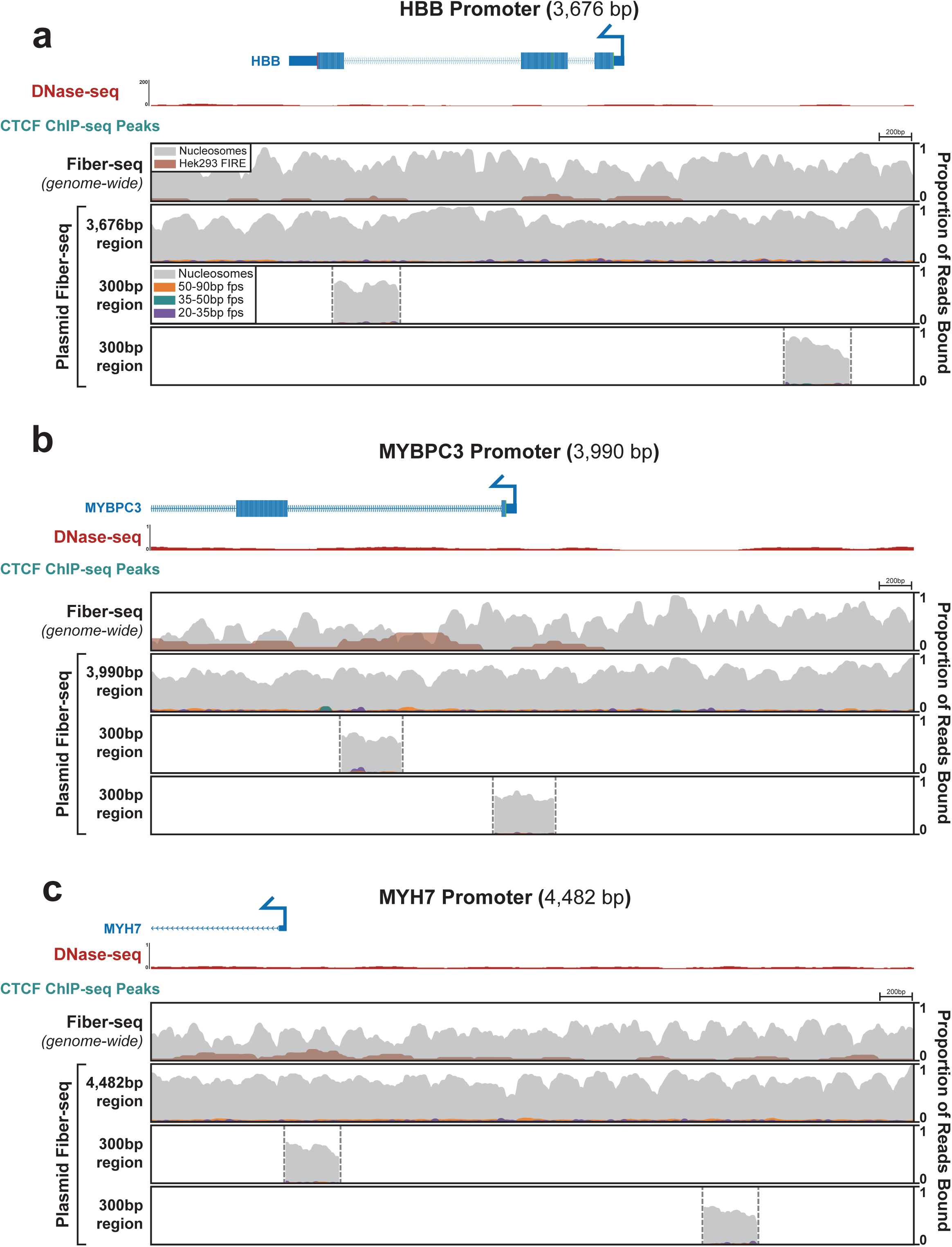
| Chromatin architecture of inactive regulatory elements Comparison of chromatin architectures of the *HBB* **(a)**, *MYBPC3* **(b)**, and *MYH7* **(c)** promoters. The top panel of each promoter shows two separate 300 bp promoter fragments centered over the Fiber-seq Inferred Regulatory Element (FIRE) peaks derived from the genomic Fiber-seq data. The middle and bottom panel show the same ∼3.6-4.5kb regions of the promoter in the plasmid and endogenous genomic context respectively. Top tracks show DNase and ChIP-seq data from HEK293 cells, along with a map of the gene body.

**Extended Data Figure 10.**
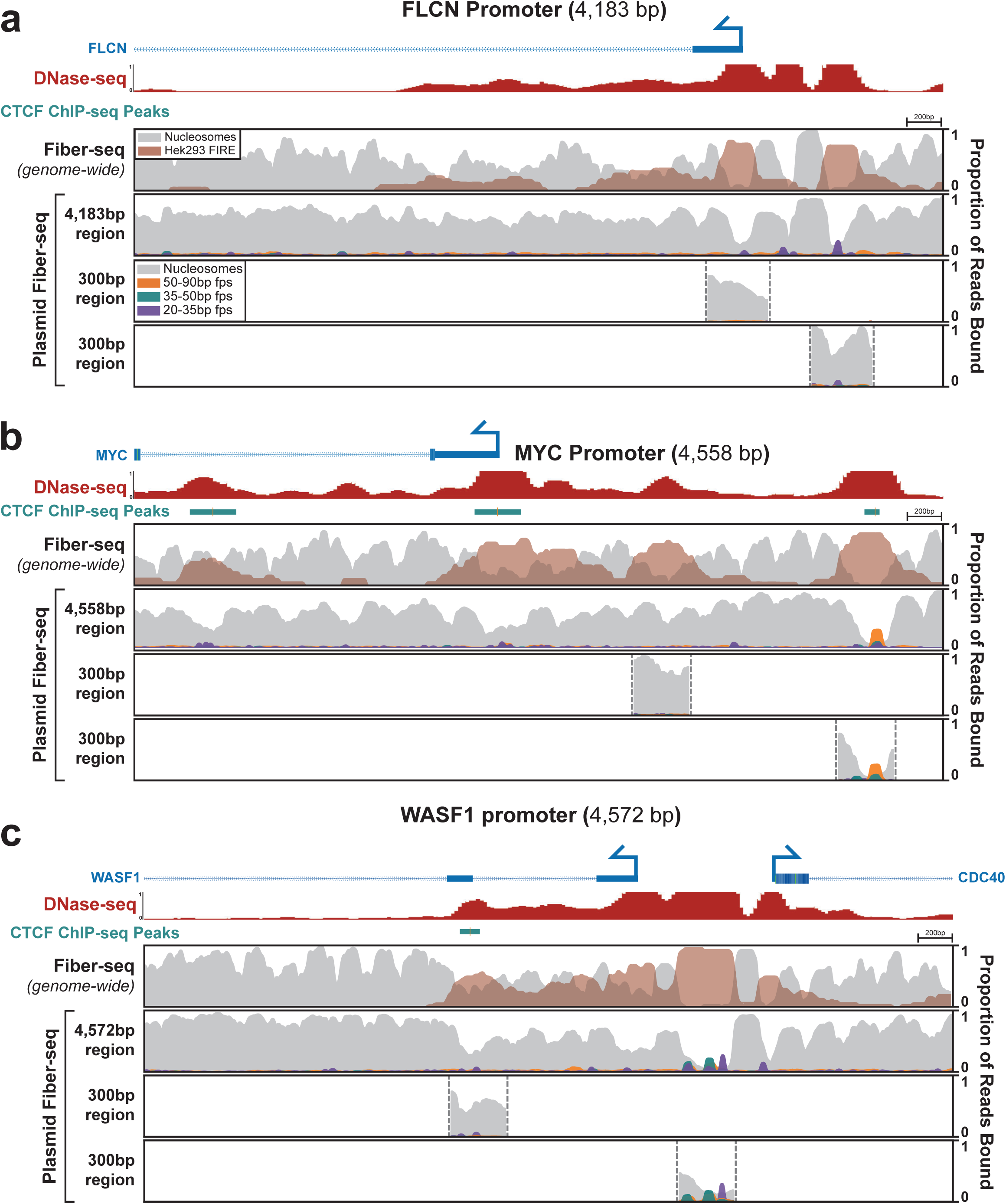
| Chromatin architecture of active regulatory elements Comparison of chromatin architectures of the *FLCN* **(a)**, *MYC* **(b)**, and *WASF1* **(c)** promoters. The top panel of each promoter shows two separate 300 bp promoter fragments centered over the Fiber-seq Inferred Regulatory Element (FIRE) peaks derived from the genomic Fiber-seq data. The middle and bottom panel show the same ∼4-4.5kb regions of the promoter in the plasmid and endogenous genomic context respectively. Top tracks show DNase and ChIP-seq data from HEK293 cells, along with a map of the gene body.

**Extended Data Figure 11.**
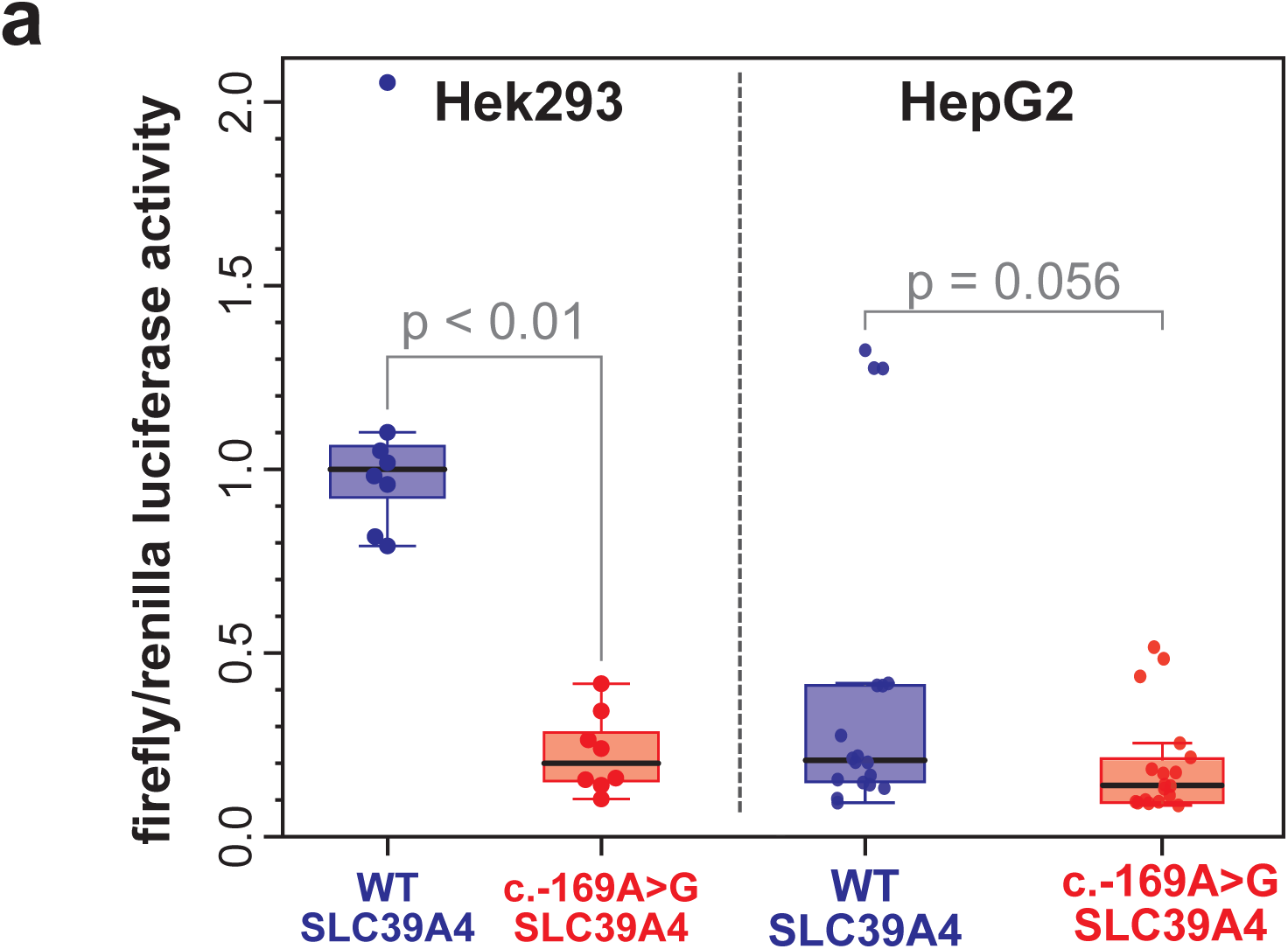
| Luciferase Assay of pSLC39A4 in Hek293 and HepG2 **(a)** Luciferase assay of the *SLC39A4* promoter region and variant promoter region in Hek293 and HepG2, demonstrating that the *SLC39A4* promoter variant clearly disrupts the promoter activity of this region in HEK293 cells, resulting in 21% promoter activity compared with the wild-type sequence. In contrast, promoter activity was weak for both the wild-type and variant promoter sequences in HepG2 cells.

**Extended Data Figure 12.**
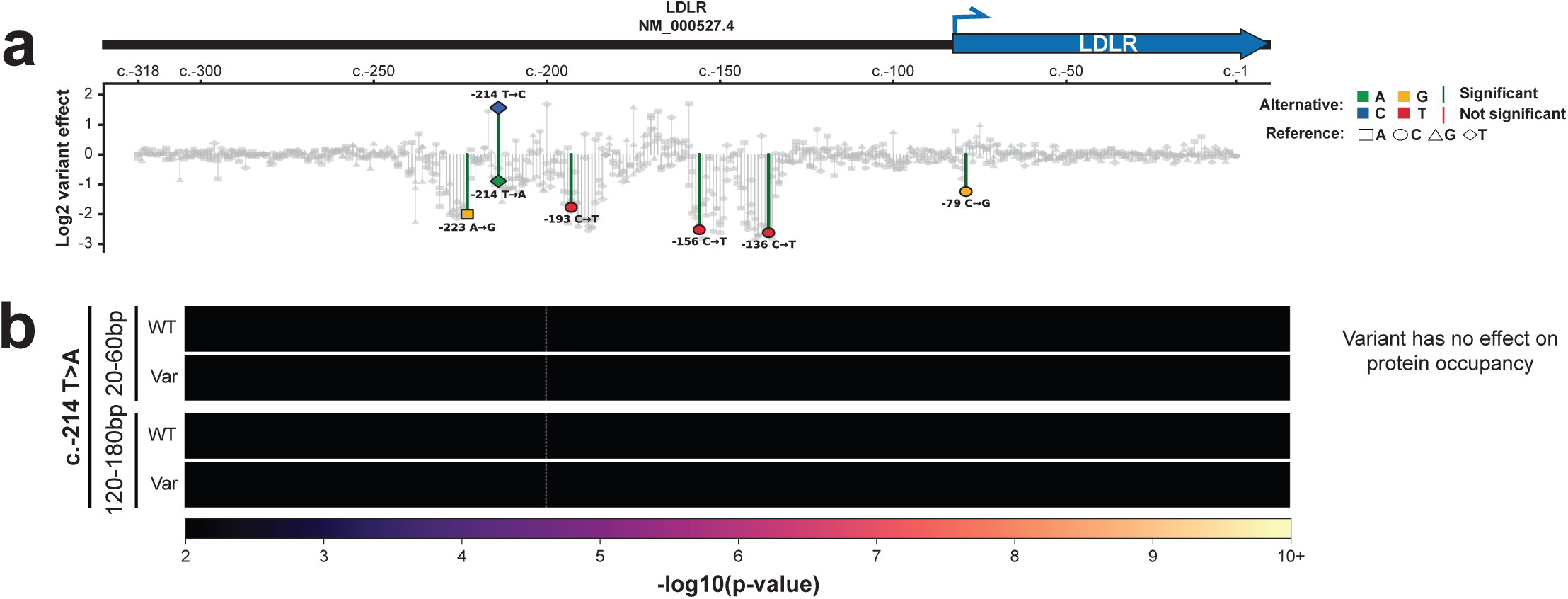
| Variant effects can be sequence specific **(a)** Log2 variant effects of all SNVs ordered by their RefSeq transcript position in NM_000527.4 of the hypercholesterolemia-associated *LDLR* promoter. Colored data points correspond to variants whose molecular mechanism of variant effect is defined in b. Significance level (red/green lines) is 10^-5^. Data for this plot is sourced from Kircher et al^32^. **(b)** Heatmaps showing the enrichment of protein footprint occupancy for both small proteins (20-60 bp) and nucleosomes (120-180 bp) along the *LDLR* promoter for both the wild type and specified variant sequence for the c.-214T>A variant. In contrast to the c.-214T>C variant that recruits an activator as shown in figure 6, this shows that a T>A variant at the same position has no effect on protein occupancy, and corresponds with a decrease in promoter activity.

**Extended Data Figure 13.**
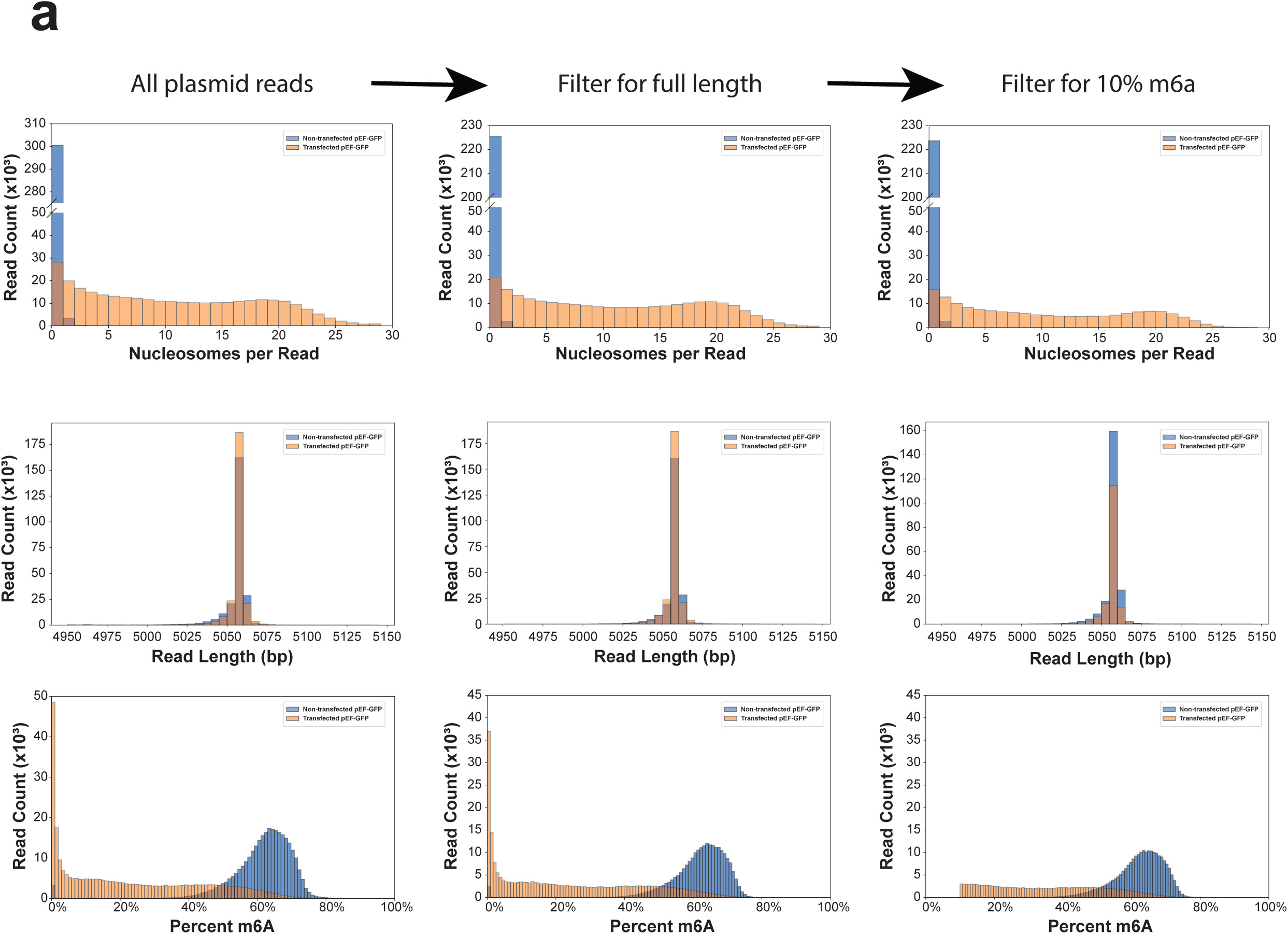
| Length and m6a filtering of plasmid Fiber-seq reads **(a)** Histograms depicting the minimal changes in the distribution of chromatin state (top), read length (middle) and methylation (bottom) when applying full read length (reference length +/- 50 bp) and methylation cutoff (>10% methylated) filters prior to analyzing plasmid Fiber-seq data.

**Extended Data Figure 14.**
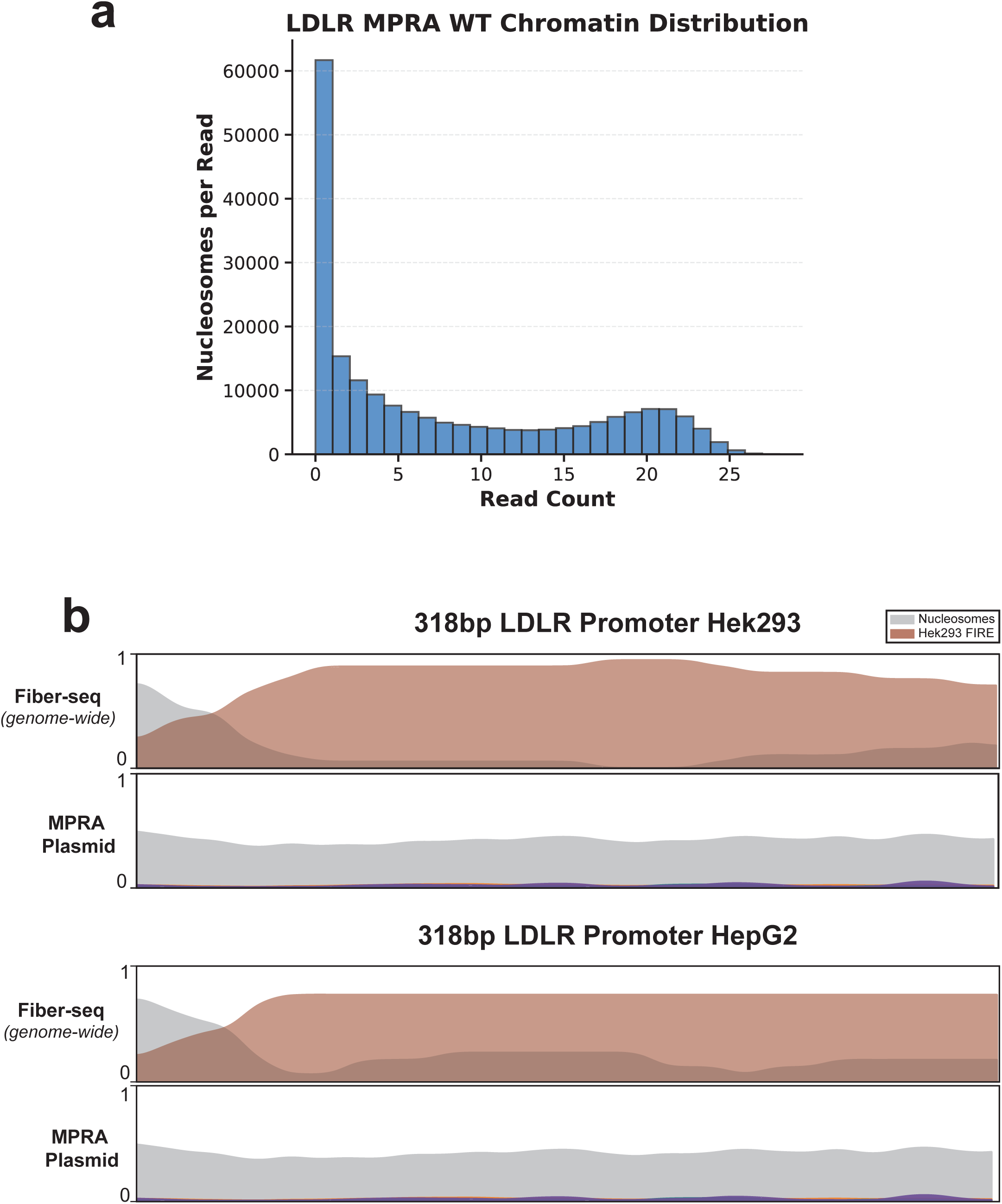
| *LDLR* Promoter Chromatin Architecture **(a)** A histogram showing the chromatin distribution of the LDLR MPRA plasmid in HEK293. **(b)** Smoothed line plots showing the chromatin architecture of the 318 bp *LDLR* promoter in the genomic (top) or plasmid (bottom) context. Comparison between HEK293 and HepG2 show highly similar architectures in both cell types. Y-axis represents the proportion of reads occupied by an element.

